# Gastrointestinal germinal center B cell depletion and reduction in IgA^+^ plasma cells in HIV-1 infection

**DOI:** 10.1101/2024.05.17.590425

**Authors:** Francesca Cossarini, Joan Shang, Azra Krek, Zainab Al-Taie, Ruixue Hou, Pablo Canales-Herrerias, Minami Tokuyama, Michael Tankelevich, Adam Tillowiz, Divya Jha, Alexandra E. Livanos, Louise Leyre, Mathieu Uzzan, Gustavo Martinez-Delgado, Matthew D. Taylor, Keshav Sharma, Arno R Bourgonje, Michael Cruz, Giorgio Ioannou, Travis Dawson, Darwin D’Souza, Seunghee Kim-Schulze, Ahmed Akm, Judith A. Aberg, Benjamin K. Chen, Douglas S. Kwon, Sacha Gnjatic, Alexandros D. Polydorides, Andrea Cerutti, Carmen Argmann, Ivan Vujkovic-Cvijin, Mayte Suarez-Fariñas, Francesca Petralia, Jeremiah J. Faith, Saurabh Mehandru

## Abstract

Gastrointestinal (GI) B cells and plasma cells (PCs) are critical to mucosal homeostasis and the host response to HIV-1 infection. Here, high resolution mapping of human B cells and PCs sampled from the colon and ileum during both viremic and suppressed HIV-1 infection identified a reduction in germinal center (GC) B cells and follicular dendritic cells (FDCs) during HIV-1 viremia. IgA^+^ PCs are the major cellular output of intestinal GCs and were significantly reduced during viremic HIV-1 infection. PC-associated transcriptional perturbations, including type I interferon signaling, persisted in antiretroviral therapy (ART)-treated individuals, suggesting ongoing disruption of the intestinal immune milieu during ART. GI humoral immune perturbations were associated with changes in the intestinal microbiome composition and systemic inflammation. These findings highlight a key immune defect in the GI mucosa due to HIV-1 viremia.

**One Sentence Summary:** Intestinal germinal center B cell reduction in HIV-1 infection linked to reduced IgA^+^ plasma cells and systemic inflammation.

## INTRODUCTION

Neutralizing antibodies (NAbs) play a critical role in host protection against invading pathogens such as HIV-1(*1*). There is a relative lack of understanding of the humoral immune dynamics in gut-associated lymphoid tissue (GALT), a site where viral replication is concentrated during HIV-1 infection (*2–4*) concomitant with a profound CD4^+^ T cell depletion within this site (*5–8*). GALT-associated HIV-1 replication is associated with seeding of HIV-1 reservoirs (*9–11*), disruption of mucosal barrier integrity and systemic inflammation (*12–17*). Therefore, a detailed examination of the humoral immune system of the GALT is important to understanding HIV-1 pathogenesis and to inform HIV-1 cure efforts.

The cellular output of GALT is dominated by secretory IgA (SIgA) that regulates barrier integrity (*18–21*) and mediates mucosal homeostasis (*22*). The generation of IgA requires B cell-T cell interactions within the highly organized microenvironment of germinal centers (GCs) (*22–24*). Within the GCs, specialized stromal cells called follicular dendritic cells (FDCs) are crucial to the maturation of the B cell response through affinity-based selection of B cells that drive the production of highly specific antibodies (*25*). B cells egress from the GC as memory B cells or as plasma cells (PCs), which then migrate to effector sites such as the intestinal lamina propria or to the bone marrow for long-term survival (*25–28*).

Intimate bidirectional crosstalk exists between SIgA and intestinal bacteria. Intestinal dysbiosis is recognized in HIV-1 infection (*29–36*), however, the impact of humoral perturbations on microbiota alterations is not well elucidated in persons with HIV-1 infection (PWH). Additionally, while a number of gut barrier-associated homeostatic mechanisms have been shown to be disrupted by HIV-1 infection (*17, 37–39*), B cells and PCs, key regulators of the gut barrier remain understudied in PWH.

## RESULTS

### Viremic HIV-1 infection is associated with a major depletion of GC B cells in the GI tract

To dissect the dynamics of intestinal B cells and PCs, single cell RNA sequencing (scRNA-seq) was performed in 9 PWH (4 people with viremic HIV (PWVH) with plasma HIV-RNA >20 copies/ml off ART and 5 people with suppressed HIV (PWSH) on ART (plasma HIV-RNA <20 copies/ml for at least 12 months) and 9 HIV-negative controls (NC) (Table S1 and S2). Immune cell analyses were conducted independently for the colon and ileum to reflect potential differences in the immune effector (colon) and inductive (ileum) sites respectively (*40*). Unsupervised cell clustering of 62365 cells with an average of 9016 Unique Molecular Identifiers (UMIs) per cell from the colon generated 23 cell clusters (**Fig. 1**, A and B, Fig. S1A and Table S3), while 43291 cells with an average of 6292 UMIs per cell from the ileum mapped to 26 cell clusters (**Fig. 1**, C and D, Fig. S1B and Table S3). Based on canonical gene expression, we identified clusters representing epithelial (*KRT8, SLC25A5*, *CA2*), endothelial (*PLVAP, PECAM*), stromal (*PDGFRB, CXCL14, CALD1, PMP22*) and immune cell lineages that included PCs (*JCHAIN, MZB1, XBP1*), B cells (*CD79A, BANK1*), CD4^+^ T cells (*CD3D, CD4, TRAC*), CD8^+^ T cells (*CD3D, CD8, TRAC*), myeloid cells (*CD14, CD68, MNDA*), innate lymphoid cells (*IL7R, KIT, IL23R*) and mast cells (*TPS, KIT)* both in the colon and ileum (**Fig. 1**, E and F and Tables S4-7). Cell type frequencies across groups are quantified in **Fig. 1**, G and H, Fig. S1, C and D and Table S8.

**Figure 1.**
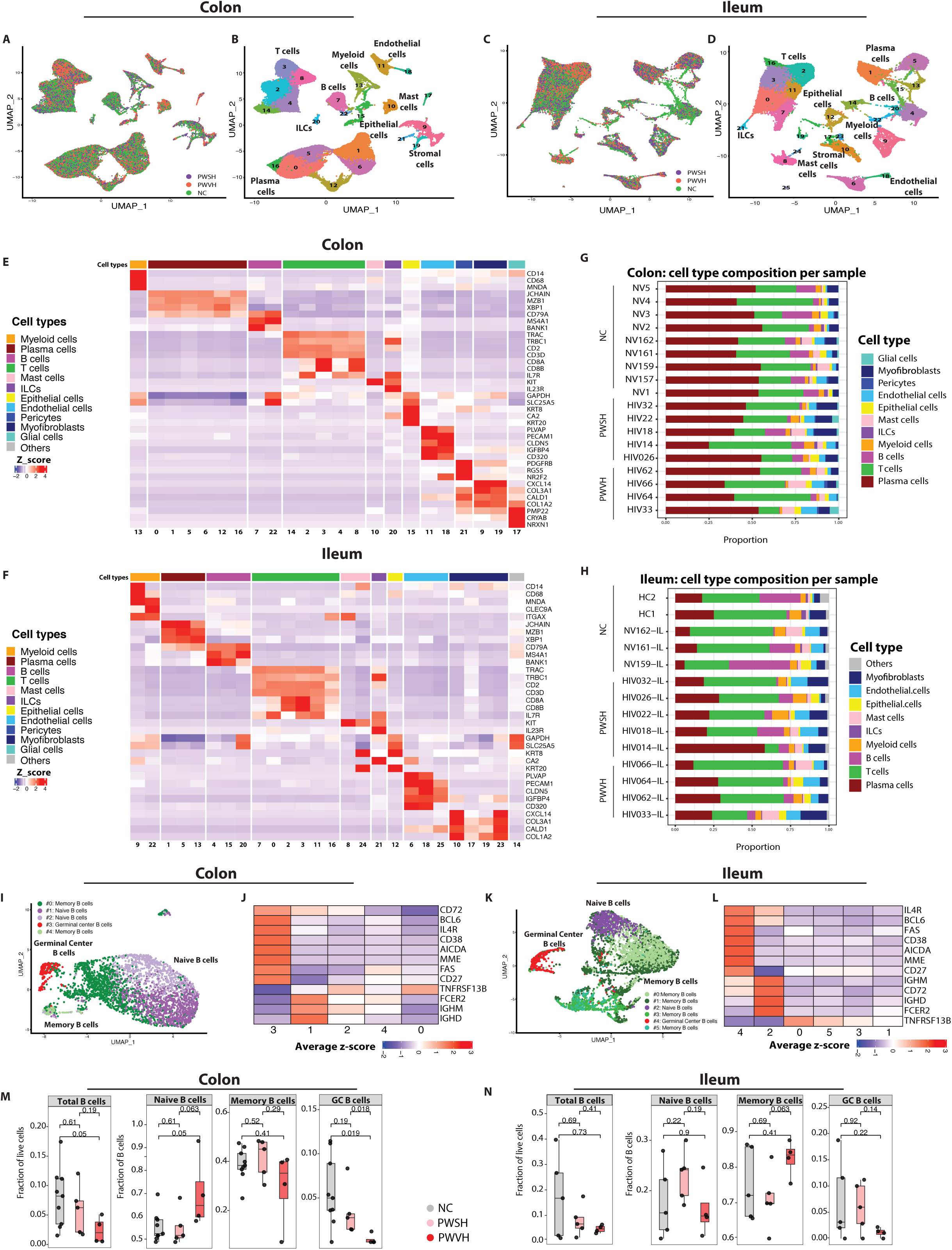
scRNA sequencing-based identification of major immune cell types in the intestinal lamina propria during HIV infection. **(A)** Uniform Manifold Approximation and Projection (UMAP) of single cell RNA sequencing (scRNA-seq) data from colonic lamina propria with representation of sample distribution across the groups (HIV negative controls (NC), n=9, green; People with suppressed HIV infection (PWSH), n=5, orange; People with viremic HIV infection (PWVH), n=4, purple). **(B)** UMAP representing cell-type annotated clusters derived from the colonic lamina propria. **(C)** UMAP of ileal lamina propria-derived samples with representation of sample distribution across the groups (NC n=5, green; PWSH, n=5, orange; PWVH, n=4, purple). **(D)** UMAP representing cell-type annotated clusters derived from the ileal lamina propria. **(E, F)** Heatmap showing the average z-score-normalized log expression of canonical cell type markers across different colonic **(E)** and ileum-derived **(F)** cell clusters. **(G, H)** Bar plots showing cell-type distribution within each individual for colon-derived **(G)** and ileum-derived **(H)** samples. **(I)** UMAPs showing re-clustered, colon**-**derived B cell clusters using scRNA-seq. **(J)** Heatmap showing the average normalized log expression z-score of canonical B cell markers across different colon-derived B cell sub-clusters. **(K)** UMAPs showing re-clustered, ileum-derived B cell clusters using scRNA-seq. **(L)** Heatmap showing the average z-score normalized log expression of canonical B cell markers across different ileum-derived B cell sub-clusters. **(M, N)** scRNA-seq data demonstrating the frequency of B cell sub-clusters derived from the colon **(M)** and ileum **(N)** across the three groups (NC, n=9 for colon and n=5 for ileum; PWSH, n=5 for colon and ileum, both; PWVH, n=4 for colon and ileum, both). Bars represents median values. P-values from Wilcoxon signed-rank sum test are reported.

Next, we re-clustered B cells using an unsupervised clustering algorithm (*41*). In the colon, 3382 B cells yielded 5 clusters, while in the ileum 5095 B cells yielded 6 clusters (**Fig. 1**, I and K and Table S9). We annotated naïve B cells (*IGHM, IGHD, FCER2, no CD27*; colon cluster #1, ileum cluster #2), memory B cells (*TNFRSF13B, CD27,* no *IGHM, IGHD*; colon clusters #0 and 2, ileum clusters #0, 1, 3 and 5) and GC B cells (*AICDA, BCL6, CD38, MME;* colon clusters #3, ileum clusters #4) (**Fig. 1**, J and L and Tables S10-13). Notably, Ig genes and gene fragments were considered and retained in the sub-clustering algorithm, as they are critical for cell state definition (specifically *IGHM and IGHD* for naïve vs. memory B cells) and otherwise did not contribute to the determination of the clusters (methods). We examined the frequency of total, naïve, memory and GC B cells across groups. GC B cells showed a -4.255 Log_2_ fold change in the colon and a -2.622 Log_2_ fold change in the ileum in PWVH compared to NC as well as a -3.517 Log_2_ fold change in the colon and a -2.380 Log_2_ fold change in the ileum compared to PWSH (**Fig. 1**, M and N and Table S14).

To validate scRNA-seq data in a larger cohort, we examined B cell subsets using multiparameter flow cytometry (FC) in 18 PWVH, 49 PWSH and 87 NC (Table S1, S2, S15, S17). Consistent with the scRNA-seq data, GC B cells (CD45^+^CD3^-^CD19^+^CD38^+^IgD^-^ Fig. S2A) showed a -1.6 Log_2_ fold change in the colon (p= 0.0027, Cohen’s d Effect Size: 0.829) and a -2.4 Log_2_ fold change in the ileum (p=0.0058, Cohen’s d Effect Size: 0.698) in PWVH compared to both PWSH and NC (**Fig. 2**, A and B, Table S17 and S18).

**Figure 2.**
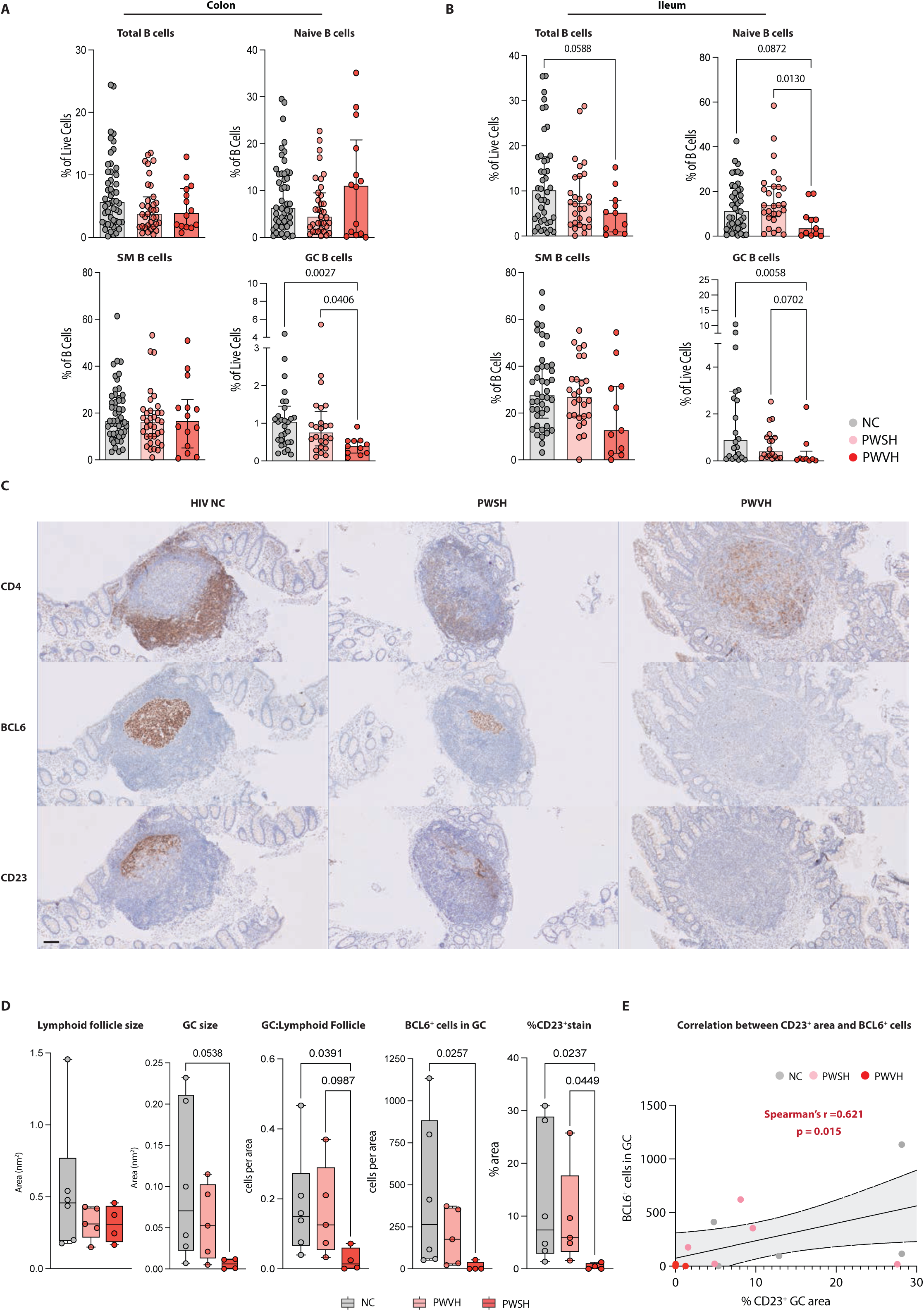
Depletion of intestinal Germinal Center B cells during viremic HIV-1 infection. (A, **B)** Flow cytometric data quantifying colon-**(A)** and ileum-derived **(B)** total B cells and B cell subsets. Comparisons between groups performed with one-way ANOVA with Dunn’s correction for multiple comparisons, p-values as indicated. **(C)** Representative immunohistochemistry (IHC) images of intestinal lymphoid follicles for each of the three groups showing staining for CD4 (top panels), and BCL6 (middle panels) and CD23 (bottom panels). Scale bar represents 100μm. **(D)** Summary data for IHC (NC n=6; PWSH, n=5; PWVH, n=4), quantifying lymphoid follicle size, GC size, the ratio between GC and lymphoid follicle size, the number of BCL6^+^ cells in each GC and the % area positive for CD23 staining within each GC. **(E)** Correlation analysis between the area with positive CD23 staining and the number of BCL6^+^ cells within each GC. Comparisons between groups were performed using Kruskal-Wallis test with Dunn’s correction for multiple comparisons. Median and interquartile range was used to plot summary data, p values <0.1 are shown as indicated (p-values > 0.1 are not shown). Correlation was obtained using Spearman’s r correlation coefficient, p values as indicated.

To further confirm our findings, we performed immunohistochemistry (IHC) on intestinal biopsy tissues where a lymphoid follicle was identified (Table S2, Table S16, Fig. S3-S5). The size of GCs was on average 16 times smaller (p=0.0538) in PWVH compared to NC, despite overall comparable lymphoid follicle size (**Fig. 2**, C and D). We used BCL6 as a marker of putative GC B cells and found a 31-fold reduction of CD4^-^BCL6^+^ cells within the GC in PWVH when compared to NC (**Fig. 2**, C and D, Table S16). Additionally, we examined FDCs, given their importance in maintaining the GC response (*25*), using CD23 staining (*42*). CD23 expression was 9-fold lower in PWVH compared to NC (**Fig. 2**, C and D, Fig. S6, Table S16) and correlated with BCL6 expression (**Fig. 2E** and S7).

Altogether, based on scRNA-seq, flow cytometry and IHC, we identify a significant loss of GC B cells in the GI tract of PWVH that is accompanied by disruption of the FDC network.

### Total PCs and IgA^+^ PCs are decreased in the colon during viremic HIV-1 infection and demonstrate an altered transcriptional profile

We next examined whether PC changes accompany the loss of GC B cells in PWVH. Using multiparametric FC we defined intestinal PCs **(Fig. S2A)** and observed a -0.9 and -1.7 Log_2_ fold change in the frequency of colonic PCs (live CD27^+^CD38^hi^cells) in PWVH, when compared to PWSH and NC respectively **(Fig. 3, A and B and Table S17-18).** Among the PC isotypes, the frequency of IgA^+^ PCs was 1.1 times lower in PWVH [median (IQR): 82 (61–91)%] compared to NC [92 (87–95)%, p=0.005, Cohen’s d Effect Size 1.251] **(Fig. 3, C and D and Table S17-18).** Concomitantly, the frequencies of IgG^+^ [6.2 (2.9-13)%] and IgM^+^ [4.7 (2.5-7.9)%] PCs were higher in PWVH when compared to NC [3.8 (2.2-7.4)% p=0.289, Cohen’s d Effect Size: 0.857 for IgG^+^ and 2.3 (1.4-3.9)%, p=0.0132, Cohen’s d Effect Size: 0.958 for IgM^+^) **(Fig. 3, C and D and Table S17-18).** PC-related changes in the ileum were less striking than those observed in the colon **(Fig. S8, A and B and Table S17).**

**Figure 3.**
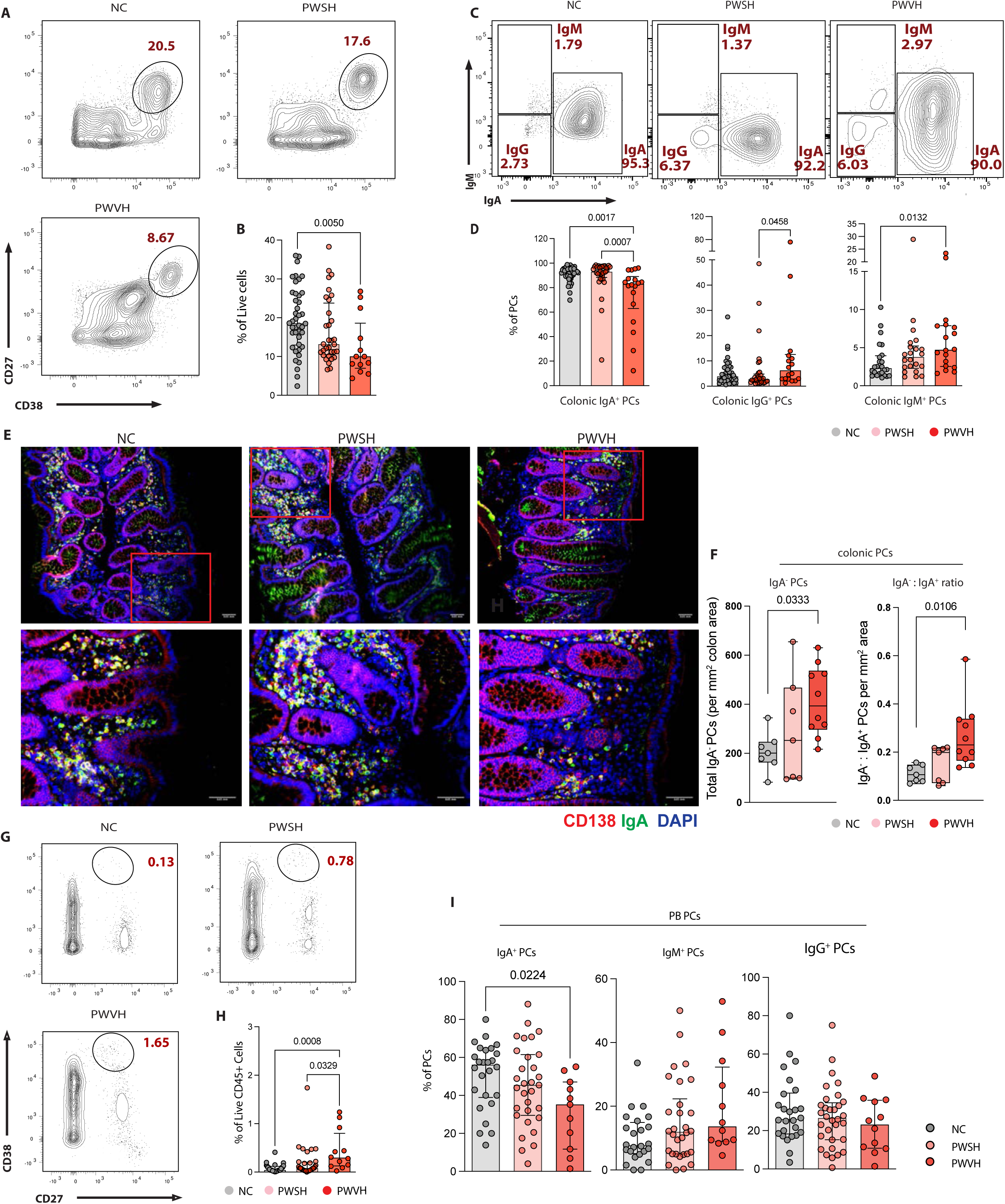
Colonic PCs are decreased during untreated HIV infection and show switch in isotype expression. **(A)** Representative flow cytometry plots of PCs (live CD27^+^CD38^hi^cells) derived from NC, n=51; PWSH, n=35; PWVH, n=16. **(B)** Summary data quantifying colonic PCs for each of the three groups. **(C)** Representative flow cytometry plots demonstrating PC isotype expression. **(D)** Summary data quantifying colonic IgA^+^, IgG^+^ and IgM^+^ PCs. **(E)** Representative immunofluorescence (IF) images showing the expression of CD138 (red), IgA (blue) and DAPI (grey). Top panel (10X magnification) with yellow insert shown in the bottom panels (20X magnification). Red cells indicate IgA^-^ PCs, blue cells indicate IgA^+^ cells and purple cells (merge) indicate IgA^+^ PCs. **(F)** Summary IF data showing total IgA^-^ PC per mm^2^ area (left panel) and ratio of IgA^-^ : IgA^+^ PCs per mm^2^ area (right panel). **(G)** Representative flow cytometry plots of peripheral blood (PB)-derived PCs (live CD45^+^CD27^+^IgD^-^CD38^hi^cells) for each of the three groups. **(H)** Summary data quantifying PB PCs. **(I)** Summary data quantifying PB-derived IgA^+^, IgM^+^ and IgG^+^ PCs. Comparisons between groups were performed using Kruskal-Wallis test with Dunn’s correction for multiple comparisons. Median and interquartile range was used to plot summary data, p values <0.1 are shown as indicated (p values > 0.1 are not shown).

Next, we quantified the absolute numbers of IgA^+^ and IgA^-^ PCs and noted that PWVH showed a 2-fold increase in IgA^-^ PCs (p=0.0333) and IgA^-^:IgA^+^ PC ratio (p=0.0106) compared to NC (**Fig. 3**, E and F and Table S19). Using the criteria defined by Landsverk et al (*43*), we characterized changes in short-lived (SL; live,CD27^+^CD38^hi^CD45^+^CD19^+^cells), long-lived (LL; live,CD27^+^CD38^hi^CD45^+^CD19^-^cells) PCs and very long-lived (VLL; live,CD27^+^CD38^hi^CD45^-^ CD19^-^cells) (Fig. S2A and Table S17) and noted similar trends as observed in the overall PCs in the colon (Fig. S8C and Table S17). In the ileum a higher frequency of LL PCs was noted for PWVH compared to NC (Fig. S8D and Table S17). Again, isotype expression across PCs subclasses showed similar trends as were observed for the overall PCs (Fig. S8E and F, Table S17).

In contrast to the intestinal mucosa, and similar to prior reports (*44, 45*), an increased frequency of PCs was observed in the peripheral blood (PB) in PWVH compared to the other two groups (**Fig. 3**, G and H, Table S17). Similar to what was observed in the colonic LP, a reduced frequency of IgA^+^ PCs was noted in the PB of PWVH compared to NC (**Fig. 3I** and Table S17).

Altogether, these data show a correlation between the perturbed GC output and the reduction in mucosal antibody-secreting PCs. Further, there is a switch in the PC isotype expression from the homeostatic IgA^+^ PCs to potentially pro-inflammatory non-IgA^+^ PC subsets. Functional studies will be needed to establish a causal relationship between reduced GC B cells and changes in PC frequency and subsets.

To further define intestinal PCs in PWH, we performed scRNA-seq on isolated mononuclear cells from the intestinal LP, and bulk RNA sequencing of fluorescence-activated cell sorted (FACS) colonic PCs. scRNA-seq-based sub-clustering of the PCs (**Fig. 4A**) revealed 12 (colon) and 8 (ileum) PC sub-clusters (Table S20). Based on immunoglobulin gene expression, PCs were grouped into IgA (colon clusters #0,1,2,3,4,5,7,8,9 and ileum clusters #1,2,5,6), IgM (colon cluster #11 and ileum clusters #0,3 [mixed IgM and IgA PCs clusters]) and IgG (colon clusters#6,10 and ileum clusters #4,7) (**Fig. 4A**, Fig. S9A and B and Tables S21-24). We investigated differentially expressed pathways for each PC isotype (FDR <0.1 **Fig. 4** B, Tables S25-27). In colonic PCs, an enrichment in interferon (IFN) response (*IRF7, ISG15, IFI6, STAT1, IRF9, IRF2, OAS2, IFNAR1, STAT2*), cell cycle regulation (*NUMA1, TUBGCP2, RAD21*), apoptosis and cell fate (*CFLAR, CASP8, FADD, TNFRSF10*), antigen presentation (*HLA-DR, TAPBP, AP1G1*) and antiviral response (*PSMA7, PSMC1, APOBEC3G*) pathways was noted, especially for IgA^+^ PCs, in PWVH. Importantly, in PWSH, differences in IFN response, cell cycle, apoptosis and cell metabolism persisted, especially in IgA^+^ and IgG^+^ PCs. Furthermore, respiratory chain pathways (*SDHA, HADHB, IDH2, FH*) were specifically upregulated in PWSH, suggesting an altered metabolic state. Pathway dysregulation was less striking in ileum PCs (Table S28).

**Figure 4.**
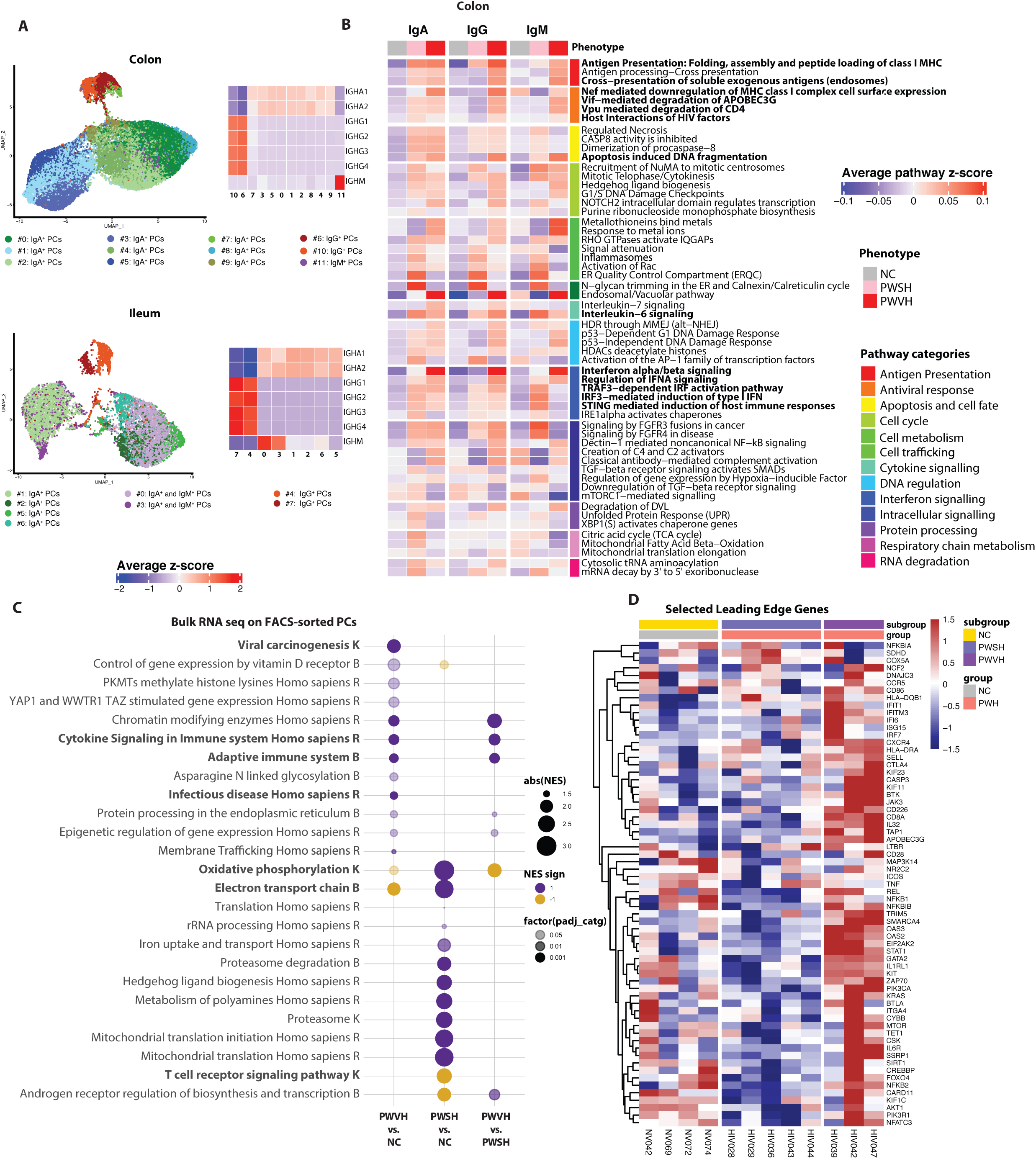
Plasma Cell transcriptional profile is altered during HIV-1 infection. **(A)** Uniform Manifold Approximation and Projections (UMAPs) (left panels) showing re-clustered, PC clusters using scRNA-seq data and heatmaps (right panels) showing the average normalized log expression z-score of Immunoglobulin genes across different colon (top panels) and ileum (bottom panels) PCs sub-clusters. **(B)** Heatmaps showing the mean pathway z-score for each colon-derived PC isotype, across the three groups (NC, n=9 for colon and n=5 for ileum; PWSH, n=5 for colon and ileum, both; PWVH n=4 for colon and ileum, both). **(C)** Bubble plot showing the Normalized Enrichment Score (NES) of selected pathways of interest derived from the top five pathways identified with Sumer analysis (methods) in FACS-sorted colonic PCs of each of the three groups (NC n=4; PWSH n=5; PWVH n=3). The bubble size represents the NES, and the color represents the direction of the change (purple = increased, yellow = decreased). Color intensity indicates statistical significance levels, with lighter colors indicating p < 0.05, darker colors for p < 0.01, and the darkest colors for p < 0.001. Pathway names ending with “K” are from the KEGG database, “B” represents Bioplanet, and “R” stands for the Reactome database. **(D)** Heatmap showing the expression of selected Leading-Edge Genes (LEGs) from pathways in panel (E) for each participant across the three groups.

To further validate these findings, we FACS-sorted colonic PCs and examined their transcriptional profile using bulk RNA-seq from 12 study participants (PWVH, n=3; PWSH, n=5; NC, n=4, Table S29). Due to the small sample size, the analysis was underpowered for detection of differentially expressed genes (at adjusted p<0.05). However, gene set enrichment analysis (GSEA) analysis leveraging different databases (i.e. KEGG, Reactome and Bioplanet) indicated dysregulation of multiple pathways (Table S30). To reduce redundancy of pathway information, we performed Sumer analysis (*46*) that uses a weighted set cover algorithm to select the fewest number of gene sets that cover all genes associated with the enriched sets (Tables S31-36). In alignment with the scRNA-seq dataset, PCs from PWVH were enriched in cell cycle (*CHAF1A*, *SETD1A, CREBBP*), IFN signaling (*IFI6, IRF7, STAT1, ISG15, OAS2, OAS3)* and innate and adaptive immune pathways (*HLA-DRA, HLA-DQB1, CD226, KIF11*) (**Fig. 4**, C and D; Tables S31-36), indicating a pro-inflammatory response in PWVH. On the other hand, PCs from PWSH showed upregulation in protein degradation (*PSMB, PSMD)* and mitochondrial metabolism pathways (*ATP6, COX5A, SDHD*) (**Fig. 4**, C and D; Tables S31-36), again demonstrating an altered metabolic state of PCs in PWSH during effective ART. Given the intrinsic methodological differences in the analysis performed via scRNA-seq and bulk RNAseq, we performed a comparative analysis of the differentially expressed pathways for each of the groups and observed an enrichment in pathways that were shared between the two methodologies across comparisons, especially in the comparisons between PWSH and NC and PWVH and PWSH (Figure S9C and Table S37).

Taken together, these data demonstrate major perturbations in the number and transcriptional profile of intestinal PCs that accompany the disrupted GC dynamics during HIV-1 viremia. Additionally, the transcriptional profile of intestinal PCs remains abnormal during ART, with persistent interferon signaling and an altered metabolic profile, further indicating that the mucosal milieu remains dysfunctional despite suppression of HIV-1 viremia.

### HIV-1-induced intestinal T cell disruption associates with PC and B cell abnormalities

Next, we studied the association of HIV-1-induced T cell changes with alterations in intestinal B cells and PCs. Using scRNA-seq, we annotated 18204 T cells in colon and 17676 T cells in ileum and identified 9 colonic T cell clusters and 10 ileal T cell clusters **(Fig. 5, A and B, Table S38).** T cells clusters mapped to CD8^+^ cytotoxic T lymphocytes (CTLs, *CD8, GZMM GZMA, GZMB, IFNG,* colon clusters #0, 2, 7 and ileum clusters #0,7), naïve and central memory T cells (*CCR7, CD27*, colon clusters #3 and ileum clusters #4) and effector memory T cells (*CD27, ICOS*, colon clusters #1, 4, 8 and ileum clusters #1, 2, 3, 6 and 9) (**Figure 5 C, D, Tables S39-42**). Additionally, we identified regulatory T cells (Tregs; *FOXP3, MAF*) in the colon (cluster #5) and ileum (cluster #8) as well as a T follicular helper cell (TFH)-like cluster (*CD40L, BCL6*) in the colon (cluster #6) and ileum (cluster #5). In PWH, the fraction of CD4^+^ T cells was decreased and the fraction of CD8^+^ T cells was increased compared to NC **(Fig. 5, E and F, Table S43).** We observed a decrease in naïve and central memory T cells and an increase in effector memory T cells in PWVH in both colon and ileum as well as a non-significant decrease in the fraction of Tregs in the colon **(Fig. 5, E and F, Table S43).**

**Figure 5.**
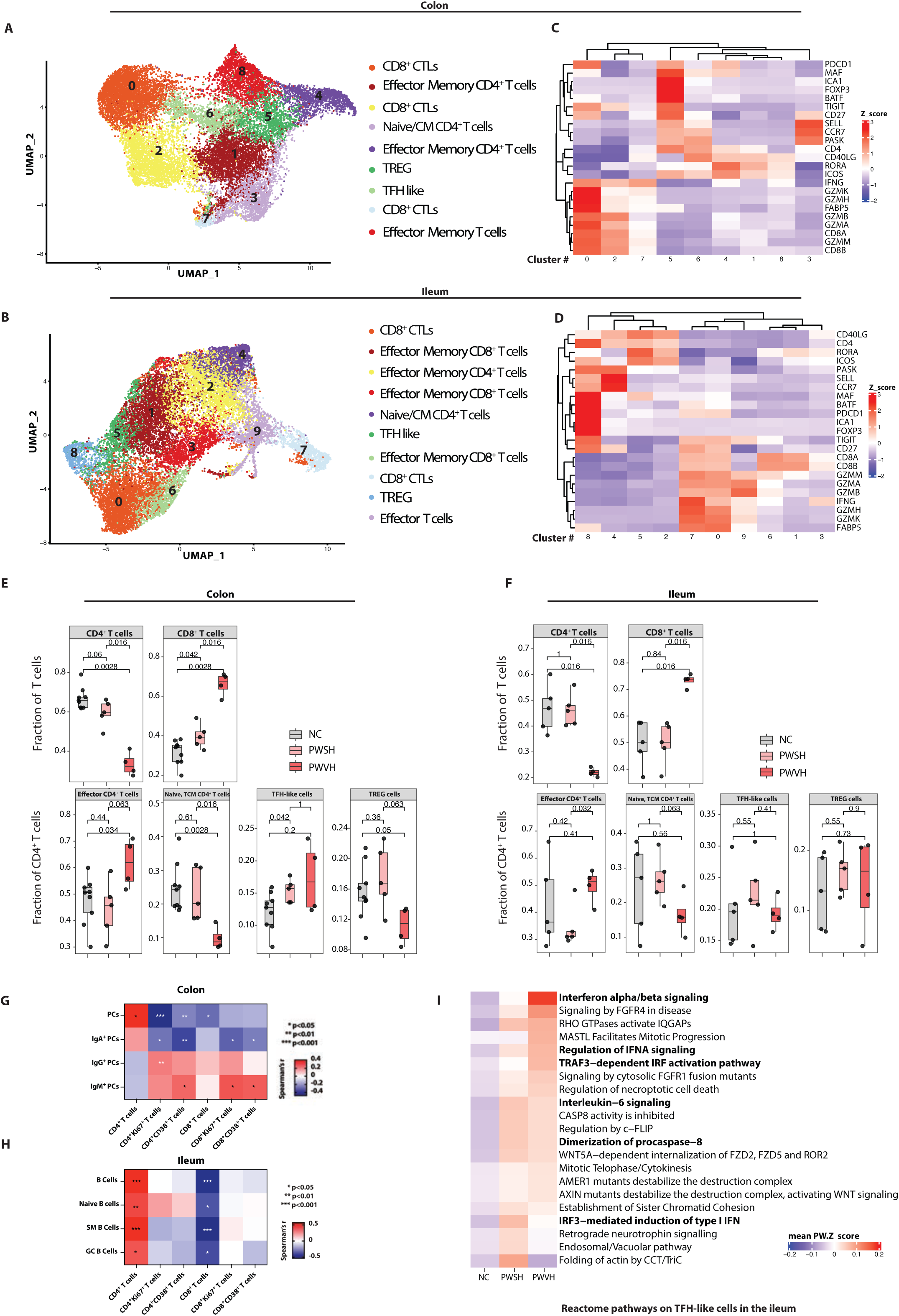
Viremic HIV-1 infection is associated with perturbations in T cell subsets including TFH-like cells. (A, **B)** Uniform Manifold Approximation and Projections (UMAPs) showing re-clustered, colon **(A)**- and ileum **(B)**-derived T cell clusters using scRNA-seq. **(C, D)** Heatmaps showing the average z-score normalized log expression of T cell related genes across different colon-**(C)** and ileum-derived **(D)** T cell subclusters. **(E, F)** Boxplots showing the scRNA-seq derived frequencies of colon-derived **(E)** and ileum-derived **(F)** CD4^+^ and CD8^+^ T cells as a fraction of all T cells (top panels), across the three groups (HIV-controls (HIV NC), n=9 for colon and n=5 for ileum; people with suppressed HIV (PWSH), n=5 for colon and ileum, both; people with viremic HIV (PWVH) n=4 for colon and ileum, both). In panels E and F, bottom boxplots show the scRNA-seq derived frequencies of colon-derived **(E)** and ileum-derived **(F)** subtypes of CD4^+^ T cells, across the three groups. In panels E and F, comparisons were performed with Wilcoxon test, p values as indicated **(G)** Correlation matrix showing relationships between frequencies of colonic PCs and PC isotypes and frequency of colonic T cell subsets. **(H)** Correlation matrix showing relationships between frequencies of ileum-derived B cells and frequency of ileal T cell subsets. Panels G and H represent derived from the flow cytometric dataset (NC, n=51; PWSH, n=35; PWVH, n=16). **(I)** Heatmap representing the mean z-score enrichment for selected pathways within the ileum-derived TFH-like T cell subcluster.

We confirmed a significant depletion of CD4^+^ T cells in the intestinal mucosa in PWVH compared to NC by FC, which did not normalize in PWSH (Fig. S10, A and B, Table S17). We also detected increased frequencies of activated (CD38^+^) and cycling (Ki67^+^) CD4^+^ and CD8^+^ T cells in PWVH compared to NC (Fig. S2C; Fig. S10 C-F, Table S17). Next, we assessed the association between intestinal T cells and intestinal PCs. We found an inverse correlation between the frequency of colonic PCs and colonic IgA^+^ PCs and activated and cycling T cells (**Fig. 5G**, Fig. S10H and I, Table S17). Conversely, a direct correlation was found between the frequency of colonic IgM^+^ (and to a lesser extent IgG^+^) PCs and activated and cycling T cell subsets (**Fig. 5G**, Fig. S10J Table S17), suggesting an association between the local inflammatory milieu and decreased PC frequency with a shift of PC isotype to non-IgA^+^ PCs. In the ileum, a direct correlation between total B cells, naïve B cells, switched memory (SM) B cells and GC B cells was observed with CD4^+^ T cells. An inverse correlation was detected between CD8^+^ T cells and total B cells, naïve B cells, SM B cells and GC B cells (**Fig. 5H**, Fig. S10K, Table S17).

Next, we examined the contribution of TFH cells in the perturbed GC dynamics. In the scRNA-seq dataset, a higher proportion of colonic CD4^+^ T cells mapping to the TFH-like cluster was noted in PWSH compared to NC. A similar, although non-significant trend was also noted for PWVH in the colon as well as for PWH in the ileum (**Fig. 5**, E and F). Since functional abnormalities within TFH cells are reported in PWH (*47, 48*), we examined the transcriptional profile of TFH cells using scRNA-seq and observed increased activation of type-I IFN, IFN-response, cell division and apoptosis pathways in PWVH in both the ileum and the colon (**Fig. 5I** and Fig. S10L, Tables S44-45). Similar, changes were seen in the TFH-like cell clusters of PWSH, suggesting that despite no detectable change in the relative frequency of TFH-like cells, the transcriptional profile of this population is altered during HIV-1 infection.

### Changes in intestinal PCs are associated with systemic inflammation

To determine whether the observed perturbations in intestinal PCs impacted the systemic immune milieu, we examined soluble and cell-associated inflammatory parameters in the circulation of the study participants. As reported previously(*49–51*), we confirmed a higher frequency of peripheral blood activated T cells (CD38^+^CD4^+^ and CD38^+^CD8^+^ T cells) in PWVH and, to a lesser extent in PWSH, compared to NC **(Fig. 6, A and B, Table S17).** PWVH also had an increased frequency of cycling (Ki67^+^CD4^+^ and Ki67^+^CD8^+^) T cells compared to HIV-NC **(Fig. 6 C and D, Table S17).** Colonic IgA^+^ PCs inversely correlated with the frequency of activated and cycling CD4^+^ and CD8^+^ T cells in circulation. Conversely, colonic IgM^+^ PCs were directly correlated with both activated and cycling CD4^+^ and CD8^+^ T cells in circulation **(Fig. 6E, Table S17).**

**Figure 6:**
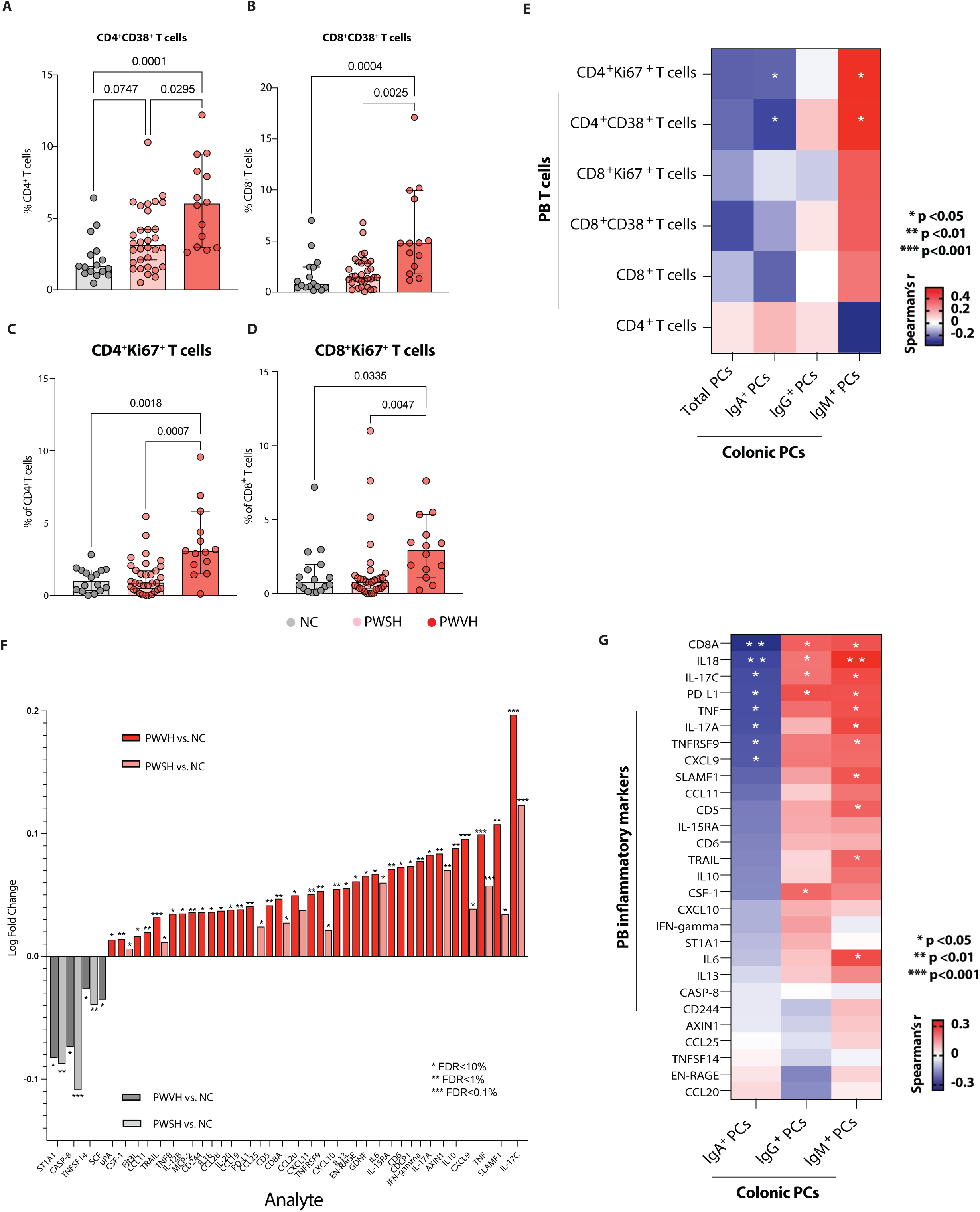
**Altered frequency and isotype expression of colonic PCs contribute to persistent systemic inflammation and immune activation during HIV infection**. **(A, B)** Frequency of peripheral blood (PB)-derived, activated (CD38^+^) CD4^+^T cells **(A)** and CD8^+^ T cells **(B)** obtained via flow cytometry, across the three groups of study participants (NC, n=33; PWSH, n=32; and PWVH, n=12). **(C, D)** Frequency of PB-derived cycling (Ki67^+^) CD4^+^ T cells **(C)** and CD8^+^ T cells **(D)** across the three groups of study participants. For panels A-D, comparisons between groups were performed using Kruskal-Wallis test with Dunn’s correction for multiple comparisons. Median and interquartile range was used to plot summary data, p values <0.1 are shown as indicated (p values > 0.1 are not shown).**(E)** Correlation matrix showing the correlation between colonic PCs and PCs isotypes and PB-derived activated and cycling T cells subsets using flow cytometry data, correlation was obtained using Spearman’s r correlation coefficient, p values as indicated **(F)** Bar plot showing systemic inflammatory markers in PWVH compared to NC (dark red) and in PWSH vs NC (light red). Dark gray bars represent systemic inflammatory markers found to be significantly higher in NC compared to PWVH and light gray bars indicate systemic inflammatory markers found to be significantly higher in NC to PWSH. Comparisons were performed by Student’s t-test with Benjamini-Hochberg correction for multiple comparisons, p values as indicated. **(G)** Correlation matrix showing the relationship between flow cytometry derived colonic PCs as well as PCs isotypes and systemic inflammatory markers noted to be elevated in People with HIV (PWH) compared to NC. Correlation was obtained using Spearman’s r correlation coefficient, p values as indicated.

We then examined a panel of soluble circulating inflammatory parameters using multiplexed proteomic assay in all study participants (n=86, (PWVH, n=13; PWSH, n=39; NC, n=34, Table S46)) where plasma samples were available. After correcting for multiple comparisons, 34 analytes were found to be significantly increased during HIV-1 infection. PWVH showed higher concentration of analytes relating to T cell activation (IL-17C, SLAMF1, IL10, IFN-γ, CD6, IL-15RA, IL6, EN-RAGE, IL-13, CCL20, CD8A, PD-L1, IL-20, CD244, TNFB, CCL11, FLT3L), myeloid cell activation (SLAMF1, TNF, CXCL9, IL6, EN-RAGE, CXCL10, TNFRSF9, CXCL11, CCL20, PD-L1, CCL19, IL-20, CD244, IL-12B), B cell activation (SLAMF1, CD6, CD5, PD-L1, IL-20) and NK cell activation (IFN-γ, IL-20, IL-18, CD244, TRAIL, FLT3L). Of these analytes, 17 remained increased in PWSH, including markers of T cell activation (IL-17C, SLMAF1, CD6, IL-15RA, IL6, CCL20, CD8A), myeloid cell activation (SLAMF1, TNF, CXCL9, CCL20, IL-20), B cell activation (SLAMF1) and NK cell activation (TRAIL) (**Fig. 6F**, Table S46). NC on the other end, had increased concentration of analytes involved in stem cell activation (SCF), T cell and dendritic cell proliferation (TNFSF14) and cell apoptosis (CASP-8). We found an inverse association of colonic IgA^+^ PCs and a direct association of IgG^+^ and IgM^+^ with most of the analytes that were increased during HIV infection (**Fig. 6G**).

Taken together, these results demonstrate perturbed T cell dynamics in the GI tract of PWH, with T cell activation associated with a decrease in colonic PCs and a shift towards non-homeostatic PC isotypes and an inverse correlation between IgA^+^ PCs and markers of systemic inflammation during HIV-1 infection.

### Discrete intestinal microbiome changes associate with distinct colonic PCs isotypes

Given the intimate, bidirectional communication of the intestinal microbiota with the mucosal immune system(*52–56*), we examined the relationship between reduced IgA^+^ PCs and the microbiota by sorting IgA-bound and unbound bacteria from stool suspensions (**methods**) in a subset of study participants (7 PWVH, 18 PWSH and 16 NC). The frequency of IgA-bound bacteria trended lower in PWVH compared to both PWSH and NC **(Fig. 7A** and **B, Table S47).** Similar findings were observed in a validation cohort (Consortium for the Evaluation and Performance of HIV Incidence Assays (CEPHIA) of the University of California, San Francisco, **methods**) where IgA binding of stool bacteria was analyzed for 5 PWVH, 13 PWSH and 12 NC **(Fig. S11A, Table S47).** To delineate bacterial species targeted by IgA-mediated host immune response, we performed IgA-seq analysis of sorted IgA-bound vs unbound bacteria in each of the three groups and calculated the log ratio of bound to unbound bacteria at genus level (*57*). The IgA log Palm score for *Actinomyces* (R=0.44, p=0.049) and *Eubacterium* (R=0.52, p=0.015) correlated with the log abundance of IgA^+^ PCs within each individual and the IgA log Palm score for *Eubacterium* inversely correlated with the log abundance of IgG^+^ PCs (**Fig. 7C** and **Table S48)** suggesting an additional homeostatic role for this genus which has previously been shown in the context of healthy gut microbiome (*58–60*).

**Figure 7:**
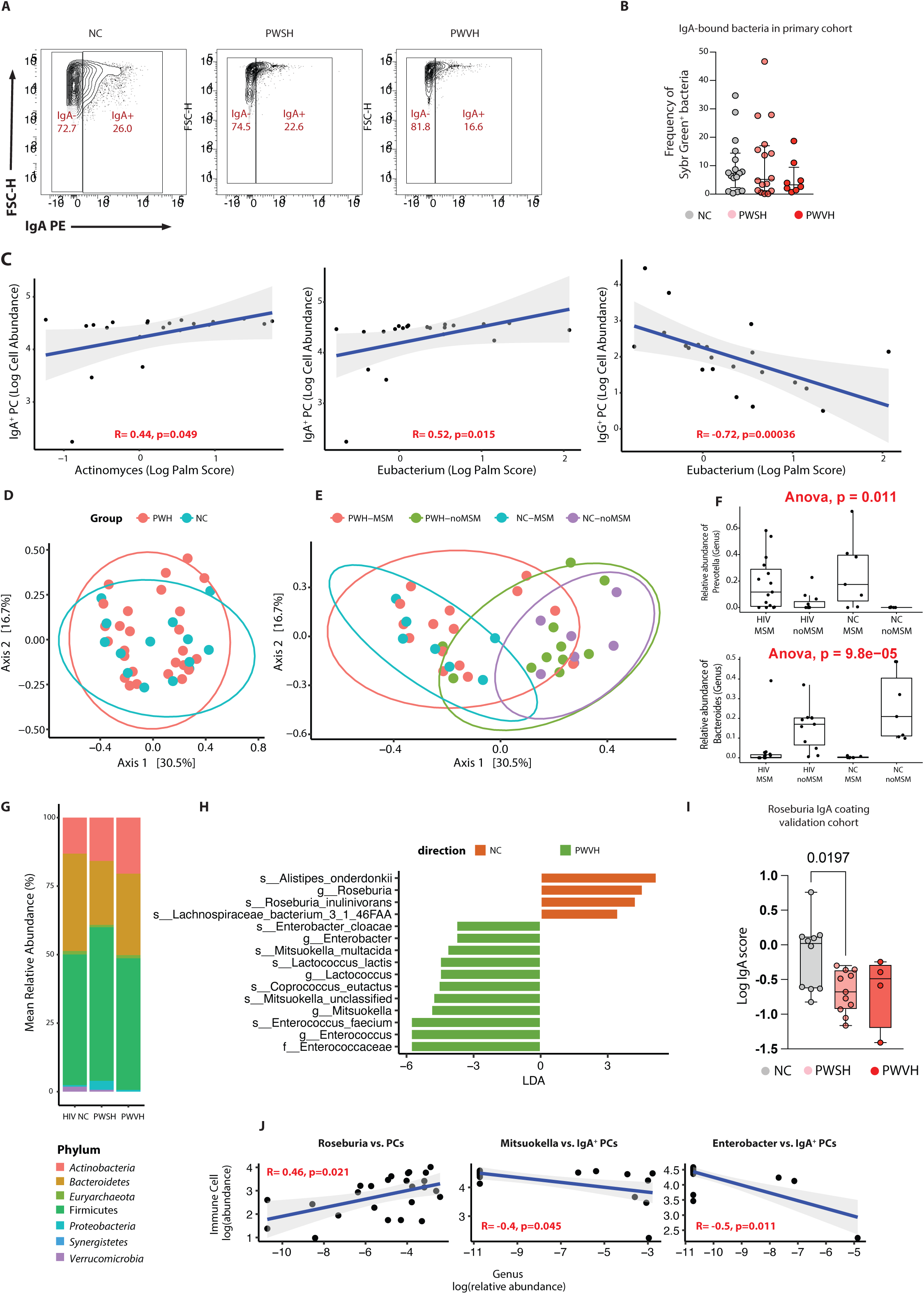
Distinct intestinal microbiota changes and targeting by secretory IgAs during HIV infection relate to altered colonic PC isotype expression. **(A)** Representative flow plots showing IgA-bound and unbound stool bacteria across the three groups (NC, PWSH and PWVH). **(B)** Summary data representing the frequency of IgA-bound bacteria across the three groups where stool samples were available for stool bacteria sorting in the primary cohort (NC, n=16; PWSH, n=18; PWVH, n=7). Bar represents median and IQR values. Statistical comparisons were performed with Mann-Whitney test, p-values as indicated. **(C)** Correlation plots between Log Palm Score and Log Cell abundance of IgA^+^ or IgG^+^ colonic PCs for bacteria where a significant correlation was found. **(D)** Principal component analysis (PCoA) plot showing Beta diversity (using Bray-Curtis distances) between PWH and NC. **(E)** PCoA plot showing Beta diversity (using Bray-Curtis distances) across PWH who are men who have sex with men (MSM) or not as well as NC who are MSM or not. **(F)** Box plots showing the relative abundance of *Prevotella* (top plot) and *Bacteriodes* (bottom plot) across PWH and NC who are MSM or not. **(G)** Barplot showing the distribution of bacterial phyla across the three groups. **(H)** Linear discriminant analysis (LDA) effect size (LEfSe) performed on all Operational Taxonomic Units (OTUs) comparing PWVH and NC. **(I)** Log IgA score for *Roseburia* in the validation cohort. Bar represents median and IQR values. Statistical comparisons performed with Mann-Whitney test, p values as indicated. **(J)** Correlation plots between the relative abundance of selected bacterial genera and the frequency of colonic PCs or IgA PCs.

To define the impact of the altered host immune landscape on the overall composition of the intestinal microbiota, we examined the stool metagenome of PWVH (n=7), PWSH (n=18) and NC (n=16). Among the 25 PWH where stool microbiota was analyzed, 15 participants identified as men who have sex with men (MSM) (**Table 1**). Since sexual practices may impact on the intestinal microbiome (*31, 61–63*), we specifically recruited HIV-negative study participants who identified as MSM (n=8, 50% of NC). Beta diversity clustering by Principal Coordinate Analysis (PCoA) revealed similar composition between PWH and NC **(Fig. 7D).** Notably, microbiota from MSM participants clustered together and were distinct from the microbiota from non-MSM participants regardless of the HIV-infection status and the microbiome distribution of HIV-infected non-MSM participants, spanned across all other subject groups **(Fig. 7E).** Consistent with previous reports (*61–63*), a higher relative abundance of *Prevotella* and lower relative abundance of *Bacteroides* was noted in MSM participants, regardless of their HIV status **(Fig. 7F).**

When examined across all three groups, the major bacterial phyla were represented by Firmicutes, Bacteroidetes and Actinobacteria, with marginal differences in their relative abundance across the three groups **(Fig. 7G).** No major differences across the three groups were noted in the relative abundance of different bacteria genera **(Fig. S11B).** Nonetheless, Linear discriminant analysis Effect Size (LEfSe), revealed enrichment of *Enterococci* and *Mitsuokella* in PWH and enrichment of *Roseburia* and *Alistipes* in NC **(Fig. 7H and Fig. S11C).** In the validation cohort, *Roseburia* was found to be less IgA-coated in PWH compared to NC as well **(Fig. 7I).**

Finally, we explored the relationship between mucosal PC frequency and the intestinal microbiota at the genus level. Frequency of colonic PCs was directly related to several microbial genera (**Fig. S11D and Table S48**) including *Roseburia* **(Fig. 7J),** which was found to be enriched in the NC in the LEfSe analysis and more IgA-coated in our validation cohort. Similarly, IgA^+^ PCs were directly associated with the relative abundance of *Bilophila, Parabacteroides, Odoribacter* and *Streptococcus* (**Fig. S11D** and **Table S49**), and inversely associated with the relative abundance of *Enterobacter* and *Mitsukella* (**Fig. 7J**), both enriched in PWVH.

Taken together these results demonstrate reduced bacterial targeting by IgAs during viremic HIV infection and alterations in the relative abundance of specific bacteria, despite the lack of major changes in the overall microbiome composition in PWH.

## DISCUSSION

NAb production is a highly coordinated process involving antigen-directed participation of TFH cells, GC B cells and FDCs (*64–67*) and GC B cell frequency relates to the magnitude of NAb titers (*68*). A majority of the identified broadly neutralizing anti-HIV-1 antibodies are highly mutated (*69*), emphasizing a pivotal role of GC B cells in the NAb response. Trama *et al.* have demonstrated that the GALT-generated responses to HIV-1 are limited and predominantly comprise microbiome-directed B cells (*70*). Additionally, Planchais *et al.* have reported that intestinal B cell-derived antibodies are largely polyreactive, with low affinity to HIV-1 envelope glycoproteins (*71*). Our report highlights a fundamental injury to the high-affinity NAb-generating apparatus in the GI tract and provides mechanistic insights regarding an impaired GALT antibody response in HIV. We posit that the loss of intestinal GC response likely contributes to a failure of virologic control, resulting in the establishment of viral sanctuaries in the intestines. This represents an HIV-specific phenomenon, as most infections and vaccinations induce a GC response (*72–76*).

HIV-1-associated GC dysfunction is likely multifactorial, including Nef-mediated perturbations of B cell function (*77*) and disruption of cognate interactions at the T cell-B cell immune synapses (*78*) as well as Vpr- and Vpu-mediated reduced B cell class switching (*79*) and antigen presentation, respectively (*80*). Chronic antigen persistence within GCs is associated with an IL-6-enriched, cytokine driven expansion of TFH cells (*81*). Yet, as shown by our data and from prior reports (*47, 48, 82*), TFH cells remain functionally altered in PWH and likely provide poor support to the GC reaction. Additionally, GC infiltration by CD8^+^ T cells and NK cells (*83*) and loss or fibrosis of the stromal network of the GALT likely contributes to altered GC dynamics in HIV-1 (*84*). Herein, we also identify a reduction of GALT FDCs, reminiscent of degenerative changes in the FDC network in peripheral lymph nodes in patients with advanced HIV-1 disease (*85, 86*). Direct infection of FDCs by HIV-1 remains unproven (*87*), while targeted killing by cytotoxic CD8^+^ T cells and NK cells, given high HIV-1 loads within FDCs, is a favored hypothesis (*88*).

IgA is the predominant antibody to emerge from intestinal GCs (*89, 90*), making it the most abundant immunoglobulin in the intestines (*91, 92*). Inflammatory stimuli, in particular T_H1_ cytokines such as IFN-γ redirect immunoglobulin class switching towards IgG (*93*). Here we describe an increase in intestinal IgG and IgM response during viremic HIV infection that is somewhat in contrast to the response induced by the majority of intestinal parasitic, bacterial and viral pathogens as IgA remains the mainstay of protection against *Giardia* (*94*), *Salmonella* (*95*), *Enterococcus* (*75*), *Vibrio cholerae* (*76*) and rotavirus infection (*96*) and vaccination (*97, 98*). Further, we observe that suppression of plasma HIV VL results in ‘normalization’ of intestinal IgA responses, consistent with a prior report by Zaunders *et al*. (*99*) but in contrast to Planchais and colleagues’ findings (*100*). Notably, in the latter study, participants who had started treatment during acute HIV infection, showed normalization of IgA^+^ B cells and PCs, suggesting that timing of initiating of ART may influence immune reconstitution (*101*).

In addition to altered PC frequency and isotypes, we have also identified altered transcriptional states of intestinal PCs in PWH, including upregulation of IFN-responses, antigen presentation and intracellular and cytokine signaling in PWVH. The transcriptome of intestinal PCs remains perturbed in PWSH even after 5 years of fully suppressive ART, with an increase in mRNA transcripts encoding for oxidative phosphorylation and mitochondrial metabolism. These data emphasize that mucosal dynamics remain perturbed even after years of apparently ‘well controlled’ HIV-1 infection.

In our data, consistent with previous reports (*61–63*), sexual practices appeared to be a driver of dysbiosis. Additionally, we have observed that the microbiome in PWVH, lacked bacterial genera and species known to be producers of butyrate (*Lachnospiraceae* and *Ruminococcaceae*), a short chain fatty acid (SCFA) important for mucosal integrity and health (*102–104*) while it was enriched for *Mitsoukella* (*30*), a bacterium associated with increased concentration of pro-inflammatory metabolites (*105–107*). Furthermore, we observed *Mitsoukella* to be inversely associated with colonic PC frequency. Conversely, other butyrate-producing bacteria of the *Odoribacter* genus were positively associated with the frequency of the homeostatic IgA^+^ PCs and inversely with the frequency of potentially pro-inflammatory IgG^+^ PCs, suggesting a contribution by *Odoribacter* to mucosal homeostasis in patients with HIV-1 infection, as in other inflammatory diseases (*108–110*). *Roseburia* a genus that is positively associated with general health (*111, 112*), was depleted in subjects with HIV-1 viremia in our cohort, consistent with prior reports (*33, 34, 113, 114*) and was directly related to the frequency of colonic PCs. In validation of our findings, samples from the CEPHIA cohort (*115*) demonstrated significantly lower IgA binding of *Roseburia* in PWH as compared to NC. Additionally, we observed a trend towards lower IgA-bound bacteria in PWVH and a higher abundance of IgA-bound *Actinomyces* and *Eubacterium*. IgA-bound *Eubacteria* were inversely correlated with the abundance of IgG^+^ PCs, highlighting the homeostatic properties of this genus (*58–60*). These data in aggregate demonstrate a bidirectional relationship between mucosal PCs and homeostatic microbiota in health that is disrupted in PWH.

Finally, we examined the associations between mucosal PCs and measures of systemic inflammation during HIV-1 infection. Consistent with prior reports (*51*), we confirmed heightened systemic inflammation and its persistence despite effective ART. The frequency of mucosal IgA^+^ PCs was inversely associated with most pro-inflammatory parameters analyzed. The opposite was true for IgG^+^ and IgM^+^ PCs, demonstrating an association between the mucosal humoral compartment and control of systemic inflammation in HIV-1 infection during ART.

As potential caveats and limitations, it is important to highlight the heterogeneity of the human GALT and PC frequencies across its length and across age (*116–118*), which could potentially impact the interpretation of our findings. The frequency of IgM^+^ PCs is lower in the present study when compared to a previous report by Magri *et al.* (*119*) that could be related to a combination of factors that include differences in biological specimens, processing protocols, variable distribution of IgM^+^ PCs across the GI tract and inter-individual variability. Finally, functional studies will be necessary to prove a causal impact of the perturbed GC dynamics and the observed alterations in PC frequency and isotype composition.

In summary, the present report details previously underappreciated disruptions in intestinal B cell and PC compartments during HIV-1 infection. Significant reduction in intestinal GC B cells during viremic infection may underlie the key changes in PC frequency. Further, alterations in the frequency, isotype expression and transcriptional profile of PCs and their association with the local and systemic inflammatory environment emphasize potential contributions of the intestinal humoral response to chronic morbidity associated with HIV even during effective ART.

## MATERIALS AND METHODS

### Study design and ethical considerations

This is a prospective observational study with the objective of studying the changes in the humoral immune responses in the GALT in PWH. Participants were recruited from the Icahn School of Medicine at Mount Sinai. Informed consent was obtained from all study participants. The study protocol was approved by the Mount Sinai institutional review board (HS#16-00512/ GCO# 16-0583). We enrolled PWVH who were either: ART-naive or within ten days of starting antiretroviral treatment; or who self-discontinued ART and had HIV RNA >1000 copies/mL at the time of enrollment. We also enrolled PWSH who had well-suppressed HIV-1 RNA (<20copies/mL) for >3months while receiving ART. Isolated viral “blips” (one time HIV RNA >20 copies/mL but <1000 copies/mL) were allowed. Additionally, we enrolled HIV-negative controls (**Table S1**). We excluded participants who were pregnant or had a concomitant intestinal infection. All data points obtained were used for analysis.

### Ileocolonoscopy, biopsy processing and lamina propria isolation

Mucosal biopsies were obtained using standard large cup biopsy forceps (n=30, limiting the effect of variability in sampling) and were transported in Roswell Park Memorial Institute medium (RPMI, GIBCO cat #11875085) on ice and processed within 2 hours of collection. Similarly, peripheral blood samples were obtained at the time of ileocolonoscopic procedure and processed within 2 hours of collection to isolate Peripheral Blood Mononuclear Cells (PBMCs) via Ficoll-based separation as previously described (*120*). All samples (including the PB) were processed fresh (to avoid the impact of cryopreservation on PC viability).

Intestinal mononuclear cells were obtained as previously described (*120*). Briefly, two 20-minutes rounds of dissociation to remove the epithelial layer [Dissociation media: 10 ml HBSS (free of calcium and magnesium) EDTA (0.5M, pH 8, Invitrogen) and HEPES (1M, Lonza)] were followed by a 40-minutes round of digestion at 37°C using gentle agitation (180 rpm) [digestion solution (10ml/sample) containing RPMI, FBS (Corning 35-010-CV), 0.005 g of Type IV collagenase (Sigma-Aldrich, cat # C5138-1G) and DNAase I (Sigma Aldrich Fine Chemicals Biosciences cat # 10104159001)]. The digested tissue was mechanically disrupted by syringe aspiration and sequential filtering through a 100-µm and a 40-µm cell strainer and washed with RPMI twice. All experiments were performed once per study participant.

### Multiparameter Flow Cytometry

Intestinal lamina propria cells and PBMCs were stained with cell type-specific antibody panels. (**Table S15)**. The cell suspension was fixed using either 2% formaldehyde (for extracellular markers staining only) or fixation/permeabilization buffer (Invitrogen # 00-5523-00) for intracellular staining. Samples were acquired using LSR Fortessa (BD). Each intestinal lamina propria cell suspension was acquired in its entirety and a minimum of 500,000 PBMCs were acquired [median (IQR) of PBMCs acquired: 927,000 (513,000–1,070,000) for the entire cohort] given the low frequencies of PCs in circulation during health. The gating strategies for annotating B cell subsets, PC subsets and activated T cell subsets are shown in Fig. S2. Data were analyzed using FlowJo v10 (Tree Star). Dead cells and doublets were excluded from all analyses. Statistical analysis was performed using Prism GraphPad software by Dotmatics. Comparison across groups was done with one-way ANOVA for non-parametric variables (Kruskal-Wallis tests) with Dunn’s correction for multiple comparisons. Median and interquartile range was used to plot summary data.

### Immunofluorescent microscopy

Immunofluorescence microscopy staining was performed on biopsies from NC (n=7), PWSH (n=7) and PWVH (n=10). Formalin-fixed, paraffin-embedded (FFPE) intestinal biopsy tissue was cut in 5µm sections. Tissue was dewaxed in xylene and rehydrated in graded alcohol and phosphate-buffered saline (PBS). Heat-induced epitope retrieval (HIER) was performed on sections submerged in target retrieval solution (Dako, S1699) in a pressure cooker. Non-specific binding was blocked with 10% goat serum for 1 hour at room temperature. Tissue was then incubated in primary antibodies (**Table S15**) diluted in 10% goat serum overnight at 4°C. Slides were then washed and incubated in secondary antibody (**Table S16**) and 4′,6-diamidino-2-phenylindole (1 μg/mL) for 1 hour at room temperature. Sections were mounted with Fluoromount-G (Electron microscopy sciences, #1798425). Controls included, no primary controls and isotype controls. Sections were imaged using a Nikon Eclipse Ni microscope and digital SLR camera (Nikon, DS-Qi2).

Analysis was done in ImageJ/Fiji®. IgA^+^ PCs (IgA^+^CD138^+^) and IgA^-^ PCs (CD138^+^IgA^-^) were quantified per area of lamina propria (median area 0.62 mm^2^, range= 0.21 mm^2^ to 1.48 mm^2^). CD138^+^ cells were only counted in the lamina propria. Statistical analysis was performed using Prism GraphPad software by Dotmatics. Frequency of cells per unit area were compared using one-way ANOVA for non-parametric variables (Kruskal-Wallis test) with Dunn’s correction for multiple comparisons. Median and interquartile range was used to plot summary data.

### Immunohistochemistry staining

152 of 154 participant samples were available for analysis. All samples were reviewed by a clinical GI pathologist, and in 15 samples, a lymphoid follicle with a GC was identified. This included 4/18 (22%) PWVH, 5 out of 49 (10%) PWSH and 6 out of 87 (6.9%) NC. All the samples where a lymphoid follicle with GC was detected were used for the analysis. Slides were baked at 70°C for 60 minutes and then loaded onto an automated stainer (Leica Bond III for CD4 staining and Dako Omnis (Dako / Agilent) for BCL6 staining. HIER was achieved with Epitope Retrieval Solution 2 for 30 minutes at 100°C (CD4 staining) Envision Flex TRS High Solution for 30 minutes at 97°C (BCL6 staining). Ready to Use Antibody reagent was added as marker and incubated for 15 min (CD4: Leica Bio systems, clone 4B12; BCL6: Dako / Agilent, Clone PG-B6p). Detection was achieved using Leica DAB Detection Kit (Bond Polymer Refine Detection) for CD4 and Envision FLEX DAB^+^ (Diaminobenzidine) (Dako Omnis) for BCL6. Slides were dehydrated and mounted. Positive and internal negative controls were employed. Lymphoid aggregates and GCs were demarcated by a pathologist on hematoxylin and eosin (H&E)-stained sections and BCL6^+^ cells were counted manually within the GCs on IHC slides. CD23 staining within the GC was measured via the *thresholder* function in QuPath®. After color deconvolution, a threshold of 0.25 was set for positive staining. The area of positive staining was calculated as % of GC area. Statistical analysis was performed using Prism GraphPad software by Dotmatics. Comparison across groups was performed by Kruskal-Wallis test with Dunn’s correction for multiple comparisons. Median and interquartile range was used to plot summary data.

### Lamina propria cell sorting and mRNA extraction

Twelve study participants (PWVH, n=3; PWSH, n=5; NC, n=4, **Table S29**) had colonic samples analyzed via bulk RNA sequencing of flow-sorted PCs. Lamina propria single cell suspension was obtained as described above, was stained with a dedicated Ab cocktail for B cell and PCs lineage (Live CD45^+^CD38^high^CD27^+^CD19^+^ cells) and sorted using BD FACS Aria II system. Sorted cells were then pelleted and RNA was isolated using RNeasy Plus Micro extraction Kit Qiagen # 74034) as per manufacturer instructions. RNA libraries were prepared using either SMART-Seq v4 Ultra Low Input Nextera XT or SureSelect XT RNA Direct v6 (**Table S29**)

### Bulk RNA sequencing analysis of flow-sorted PCs

The mean (std deviation) depth of sequencing was 76.2 ± 12.5 million, for a total of 990.6 million reads for the bulk RNA-seq dataset. Pre-processing was done as previously reported (*121*). Features with fewer than 4 normalized counts across all samples were filtered out. The counts were normalized by using voom (*122*). Normalized log counts per million and associated precision weights were then entered into the limma analysis pipeline (*123–125*). RNA-seq data were adjusted for RIN score using a linear model (*124–126*). Expression profiles were modeled using linear mixed-effects models including age and group variable (NC, PWSH, PWVH) as fixed effects, and a random intercept for each subject. Gene Set Enrichment Analysis (GSEA) was performed using clusterProfiler R package to identify the enriched terms (p.adjusted<0.05) in Kyoto Encyclopedia of Genes and Genomes (KEGG), Reactome and Bioplanet, for the comparison of the three groups. GSEA resulted in numerous enrichments indicating dysregulation of a large number of pathways: 659 for suppressed vs control; 320 for PWVH vs NC; 640 for PWVH vs PWSH **(Table S30)**. To summarize and consolidate GSEA results, pathways with adjusted p-values <0.05 were selected from each database into the sumer R package (*46*), which was used to select the most representative pathways. This process reduced the number of pathway terms organized to various clusters as follows: for PWVH vs NC 5 clusters of 87 pathways, for PWSH vs NC 8 clusters of 143 pathways and for PWVH vs PWSH 4 clusters of 67 pathways **(Tables S31-36).** The top 5 pathways (by adjusted p-value) per cluster were sub-selected for further analysis. Gene set variance analysis (GSVA) (Z-score method) was performed on the HIV gene expression data (RIN adjusted) to generate a pathway score with the core enriched genes for each pathway selected by the Sumer package. Bubble plot of Net Expression Score (NES) of selected pathway scores across groups was constructed. The expression (RIN adjusted) levels of selected leading edge genes (LEGs) across each sample of each group were displayed in a summary heatmap. The color of the heatmap in each row is based on the Z score values calculated by centering and scaling the data by standard deviation using the formula (X ∼m)/SD, where X is individual values, and m is the mean of the row.

### Lamina propria processing for single cell RNA sequencing

Samples from eighteen study participants (PWVH, n=4; PWSH, n=5; NC, n=9, **Table S2**) underwent scRNA-seq analysis. LP mononuclear cell suspension was obtained from freshly processed intestinal biopsies as detailed above and dead cells were removed using Dead Cell Removal by magnetic binding (Miltenyi Biotec# 130-090-101) as per manufacturer instructions. Cells were loaded on one lane of the 10X Genomics NextGem 3’v3.1 assay as per the manufacturer’s protocol with a targeted cell recovery of 8,000 cells. Gene expression libraries were prepared as per 10x Genomics protocol(*127*). Libraries were quantified via Agilent 2100 hsDNA Bioanalyzer and KAPA library quantification kit (Roche Cat# 0796014001) and sequenced at a targeted depth of 25,000 reads per cell. Libraries were pooled in equimolar concentration and sequenced on the Illumina NovaSeq 100 cycle kit with run parameters set to 28x8x0x60 (R1xi7xi5xR2).

### Single-cell RNA data processing and analysis

#### Filtering, normalization, batch correction

Only cells with total UMI counts greater than 1,000 and mitochondria gene fraction less than 10% were considered for the analysis. The analysis of sc-RNA data including data normalization and batch effect correction was performed via the Seurat package (*41, 128*). First, each cell was normalized independently using function *NormalizeData* which divides feature counts for each cell by the total counts for that cell. Data was then natural log transformed. Next, variable features were found using function FindVariableFeatures(*41, 128*).This function first fits a line to the relationship of log(variance) and log(mean) using local polynomial regression; then, standardizes the feature values using the observed mean and expected variance (given by the fitted line). Feature variance was then calculated based on the standardized values. Finally, features which were repeatedly variable across datasets were selected for integration via function *SelectIntegrationFeatures*(*41, 128*). Anchors were then identified using the *FindIntegrationAnchors* function, which takes a list of Seurat objects as input, and use these anchors to integrate different datasets together via the *IntegrateData* function(*41, 128*). Dimensionality reduction was performed using reciprocal principal component (PC) analysis. For visualization, the dimensionality of each dataset was further reduced using Uniform Manifold Approximation and Projection (UMAP) implemented with Seurat functions *RunUMAP*. The PCs used to calculate the embedding were as the same as those used for clustering.

#### Clustering

Clustering was performed via function *Find.Cluster* in Seurat, with resolution parameter r=0.5. We identified cluster-specific markers using function *FindMarkers* in *Seurat* (*41, 128*). Markers were ranked based on area under the receiver operating characteristic curve (AUC). Based on the cluster-specific markers and leveraging existing signatures from the literature (*129–134*), clusters were annotated into 9 cell-types for the ileum and the colon. As clustering of scRNA-seq data cells may be potentially influenced by the use of BCR and TCR gene segments, we verified the similarity of the sub-clustering structure when removing segments of BCR and TCR genes that undergo V(D)J recombination (specifically IG[HKL]V, IG[KL]J and IG[KL]C and TR[ABDG][VJC]). We chose to retain the IGH[ADEGM] genes as they are relevant in determining cell identity and function (*i.e.* naïve and memory B cells). With this approach, 84 - 97% of individual cell pairs retained the same co-classification status (quantified by the Rand Index, RI) (*135*), corresponding to the Adjusted Rand Index (ARI) of 0.61-0.89 (ARI ranges between -1 and 1, with 0 being the expected value for unrelated classifications). We thus elected to retain BCR and TCR gene segments in our sub-clustering of B cells, PCs and T cells.

#### Cell-type signatures

We considered the lineage cell-type categories to obtain gene-signature from the single-cell RNA data. Mitochondrial and ribosomal protein genes were removed in the final list of genes since they were found expressed in multiple cell-types(*41*).

#### Re-clustering of B cells, PCs and T cells

B-cells, PCs and T cells were re-clustered as smaller individual subsets using the same nearest neighborhood clustering algorithm (*41*). The resolution parameters used in re-clustering were 0.5 for PCs and T cells and 0.4 for B cells. After identifying markers for each of the subclusters we could assign them to specific subtypes of each cell type: naïve, B cells, memory B cells, GC B cells; and IgA, IgM and IgG isotypes for PCs.

#### Quantification of cell-type fractions

Cell types from primary clusters were quantified as a fraction of total cells in each sample. Cell types from subclusters were quantified as a fraction of the corresponding primary cell type in each sample unless otherwise specified. Comparisons across groups were performed using Mann-Whitney U test.

#### Pathway enrichment analysis

For each cell-type compartment (i.e., B cells and PCs), the log fold-change was computed between the average gene expression in cells derived from PWVH, PWSH and NC. Enrichment scores and adjusted p-values (FDR) were derived via gene set enrichment analysis (*136*) for PWVH vs NC and PWSH vs NC separately. Specifically, genes were ordered in decreasing order of fold-change and a Kolmogorov-Smirnov statistics was utilized to find pathways enriched at the top of the list. A permutation-based strategy was utilized to derive p-values and FDR for different pathways(*136*). Molecular pathway gene sets used in this analysis were from the Reactome database(*137*). Importantly, the transcriptome of PCs is enriched in Ig genes (*138, 139*), and accordingly, in our dataset Ig encoding genes comprised 58% ± 13% of all UMIs in PCs in colon, and 57% ± 17% in ileum, with a mean (SD) UMI number per cell for PC’s of 15195 ± 11807 in the colon and 17264 ± 12189 in the ileum.

### Stool metagenomics

Stool metagenomics was performed on all the participants for whom stools were available (7 PWVH, 18 PWSH and 16 NC, **Table S2**). Stool samples were collected from study participants before or at least two weeks after colonoscopy. Fresh samples were frozen at -80°C and mechanically pulverized. Fecal DNA for metagenomic sequencing was extracted as previously described (*140*). DNA was quantified using the Broad Range or High Sensitivity Quant-IT dsDNA Assay kit (Thermo Fisher,Q32853 and Q33130) with a BioTek Synergy HTX Multi-Mode Reader. Using NEBNext Ultra II DNA Library Prep Kit, metagenomic libraries were prepared. DNA that was fragmented by Diagenode Bioruptor Pico solictor were subjected to end repair, Illumina adaptor ligation, and purification with Beckman Coulter SPRI beads. NEBNext Ultra Q5 Master Mix was utilized to amplify ligated product using custom i5 and i7 index primer and Beckman Coulter AMPure XP beads quantified the final product. Samples were sequenced with an Illumina HiSeq (paired-end 150 bp).

### Computational analysis of metagenomic profiles

MetaPhlAn2 was used for whole-metagenome shotgun sequencing (*141*). Any sample with reads less than 100,000 was filtered out. Uncharacterized phylae and taxa that did not meet the minimum prevalence threshold of 3% of all samples were also filtered out. LEfSe was implemented to estimate bacterial candidates associated with study participants groups with a logarithmic LDA cut off ≥2.0 (*142*), using the R package *microbial.* Beta diversity (using Bray-Curtis distances) was calculated and visualized using with a principal coordinate analysis (PCoA) plot. Distance and diversity index calculations were performed using *phyloseq* (*143*). Spearman correlation was performed on bacterial genus relative abundances and immune cell abundances. Given the exploratory nature of this study, reported p-values are unadjusted.

### IgA-bound and unbound stool bacteria sorting

Fifty mg of frozen stool samples were resuspended in PBS (10x stool weight) and pelleted. The supernatant was filtered through 40 nm cell strainer. 100μL of suspension was blocked for non-specific staining using anti-mouse serum and stained for IgA (PE Mylteny Biotec, Cat#130-113-476), washed thrice and stained for DNA (SyborGreen Applied Biosciences). IgA-bound (SyborGreen^+^, IgA^+^) and IgA-unbound (SyborGreen^+^, IgA^-^) stool bacteria were sorted, pelleted, and DNA was extracted using QIAamp UCP Pathogen Mini Kit (Qiagen Cat# 50214). Staining was directed towards total IgAs and not specifically towards SIgAs, potentially capturing infiltration from monomeric serum IgAs in the stools. Comparison across groups was done with one-way ANOVA for non-parametric variables (Kruskal-Wallis tests) with Dunn’s correction for multiple comparisons. Median and interquartile range was used to plot summary data.

### 16S rRNA sequencing and IgA-seq

For DNA extraction, we utilized phenol:chloroform method on 50-200 mg of fecal specimen as previously described (*144*). We adjusted DNA templates to a concentration of 2 ng/mL. Following this, the V4 variable region of the 16S rRNA gene underwent PCR amplification using indexed primers, adopting the methodology outlined in (*145*). The resultant 16S rRNA V4 amplicons were merged and subsequently purified using AMPure XP beads (Beckman Coulter, A63880) at a bead-to-PCR reaction ratio of 1:1. Gel electrophoresis was employed to confirm the accurate amplicon size and to ensure the non-presence of primer dimers. The consolidated samples were then sequenced on an Illumina MiSeq platform, producing paired-end reads of 250bp each. To process and denoise paired-end sequence data, the DADA2 algorithm available within the QIIME2 pipeline was used. The Greengenes sequences database was used as a reference for pair-end reads merge and alignment.

We removed samples with less than 1000 reads. Ambiguously annotated phylum-level taxonomic features (e.g. “NA”) and low prevalence ASVs were removed. Additionally, ASVs less than 0.01% relative abundance in both IgA bound and unbound dataset were filtered out. A pseudocount of minimum (relative abundance of genus)/2 was computed to entries with exact 0 relative abundance entries in ASV tables. Log Palm score was calculated (*57*) to quantify IgA binding. Spearman correlation was calculated on immune cell abundances and log Palm score of bacterial genus.

A validation cohort (Consortium for the Evaluation and Performance of HIV Incidence Assays (CEPHIA) of the University of California, San Francisco) was also included to confirm the IgA-seq results. Samples were processed as previously described (*115*). To obtain genus-level IgA scores for all taxa, the DADA2 pipeline was applied and taxonomy was assigned on resulting amplicon sequence variants (ASV) using the SILVA database (v138.1, nr99). ASVs belonging to the same genus were summed, and a pseudocount of 1 was applied. The IgA score was computed as a natural log of the ratio of each genus in the IgA-positive fraction over the IgA-negative fraction.

### Multiplexed Proteomic Assay (Olink)

Circulating inflammatory cytokines were quantified in all the study participants where plasma was available (total= 86, PWVH, n=13; PWSH, n=39; NC, n=34, **Table S46**) using a multiplexed proteomic inflammation panel (Olink^®^) that simultaneously quantifies 92 inflammation-related proteins. The concentration of each biomarker was compared between PWVH and NC as well as between PWSH and NC using Student’s *t* test. The Benjamini-Hochberg procedure for multiple comparisons was used to adjust the resulting p-value. Correlation with LP cell type frequencies and circulating inflammation-related proteins was obtained by Spearman’s rho correlation coefficient.

## Acknowledgements

We acknowledge the support of the Human Immune Monitoring Core, the Flow Cytometry Core and the computational and data resources and staff expertise provided by Scientific Computing at the Icahn School of Medicine at Mount Sinai. We thank the Consortium for the Evaluation and Performance of HIV Incidence Assays (CEPHIA).

## Author contributions

The authors contributed as follows: The study was conceived and designed by FC and SM. Study participants were enrolled by FC, MTa and SM. Data was generated by FC, JS, AKr, ZAT, RH, PCH, MTo, AT, DJ, AEL, LL, MU, GMD, MTay, KS, GI, MC, TD, DDS, AK, ADP CA, MSF, FP, JJF, IVC, and SM. Analysis and interpretation was data was done by FC, JS, AKr, ZAT, RH, PCH, MTo, MTa, AT, DJ, AEL, LL, MU, ARB, MC, GI, DDS, SKS, JAA, BKC, DSK, SG, ADP, AC, CA, IVC, MSF, FP, JF and SM. Statistical analysis was done by FC, JS, AKr, ZAT, RH, PCH, MTo, MTa, ARB, GI, DDS, SG, ADP, CA, IVC, MSF, FP, JF and SM. The manuscript was drafted by FC, JS, AKr, ZAT, PCH, GI, ADP, CA, IVC, MSF, FP, JF and SM. Manuscript was revised by FC, JS, AKr, ZAT, PCH, DJ, AEL, ARB, JAA, BKC, DSK, SG, ADP, AC, CA, IVC, MSF, FP, JF and SM. The project was supervised by FC and SM.

## Data and Materials Availability

Sequencing data for bulkRNA-seq and scRNA-seq have been deposited to the GEO database (GSE272807). All source data need to evaluate the conclusions are provided in supplementary materials or in the main manuscript. Materials used to conduct this study will be made available upon request.

## References

1. D. Corti, A. Lanzavecchia, Broadly neutralizing antiviral antibodies. Annu Rev Immunol 31, 705–742 (2013).

2. M. Guadalupe, E. Reay, S. Sankaran, T. Prindiville, J. Flamm, A. McNeil, S. Dandekar, Severe CD4+ T-cell depletion in gut lymphoid tissue during primary human immunodeficiency virus type 1 infection and substantial delay in restoration following highly active antiretroviral therapy. J Virol 77, 11708–11717 (2003).

3. Q. Li, L. Duan, J. D. Estes, Z. M. Ma, T. Rourke, Y. Wang, C. Reilly, J. Carlis, C. J. Miller, A. T. Haase, Peak SIV replication in resting memory CD4+ T cells depletes gut lamina propria CD4+ T cells. Nature 434, 1148–1152 (2005).

4. R. S. Veazey, M. DeMaria, L. V. Chalifoux, D. E. Shvetz, D. R. Pauley, H. L. Knight, M. Rosenzweig, R. P. Johnson, R. C. Desrosiers, A. A. Lackner, Gastrointestinal tract as a major site of CD4+ T cell depletion and viral replication in SIV infection. Science 280, 427–431 (1998).

5. J. M. Brenchley, B. J. Hill, D. R. Ambrozak, D. A. Price, F. J. Guenaga, J. P. Casazza, J. Kuruppu, J. Yazdani, S. A. Migueles, M. Connors, M. Roederer, D. C. Douek, R. A. Koup, T-cell subsets that harbor human immunodeficiency virus (HIV) in vivo: implications for HIV pathogenesis. J Virol 78, 1160–1168 (2004).

6. J. J. Mattapallil, D. C. Douek, B. Hill, Y. Nishimura, M. Martin, M. Roederer, Massive infection and loss of memory CD4+ T cells in multiple tissues during acute SIV infection. Nature 434, 1093–1097 (2005).

7. S. Mehandru, M. A. Poles, K. Tenner-Racz, A. Horowitz, A. Hurley, C. Hogan, D. Boden, P. Racz, M. Markowitz, Primary HIV-1 infection is associated with preferential depletion of CD4+ T lymphocytes from effector sites in the gastrointestinal tract. J Exp Med 200, 761–770 (2004).

8. S. Mehandru, M. A. Poles, K. Tenner-Racz, P. Jean-Pierre, V. Manuelli, P. Lopez, A. Shet, A. Low, H. Mohri, D. Boden, P. Racz, M. Markowitz, Lack of mucosal immune reconstitution during prolonged treatment of acute and early HIV-1 infection. PLoS Med 3, e484 (2006).

9. A. Chaillon, S. Gianella, S. Dellicour, S. A. Rawlings, T. E. Schlub, M. F. De Oliveira, C. Ignacio, M. Porrachia, B. Vrancken, D. M. Smith, HIV persists throughout deep tissues with repopulation from multiple anatomical sources. J Clin Invest 130, 1699–1712 (2020).

10. J. D. Estes, C. Kityo, F. Ssali, L. Swainson, K. N. Makamdop, G. Q. Del Prete, S. G. Deeks, P. A. Luciw, J. G. Chipman, G. J. Beilman, T. Hoskuldsson, A. Khoruts, J. Anderson, C. Deleage, J. Jasurda, T. E. Schmidt, M. Hafertepe, S. P. Callisto, H. Pearson, …, T. W. Schacker, Defining total-body AIDS-virus burden with implications for curative strategies. Nat Med 23, 1271–1276 (2017).

11. S. A. Yukl, S. Gianella, E. Sinclair, L. Epling, Q. Li, L. Duan, A. L. Choi, V. Girling, T. Ho, P. Li, K. Fujimoto, H. Lampiris, C. B. Hare, M. Pandori, A. T. Haase, H. F. Gunthard, M. Fischer, A. K. Shergill, K. McQuaid, …, J. K. Wong, Differences in HIV burden and immune activation within the gut of HIV-positive patients receiving suppressive antiretroviral therapy. J Infect Dis 202, 1553–1561 (2010).

12. J. M. Brenchley, D. A. Price, T. W. Schacker, T. E. Asher, G. Silvestri, S. Rao, Z. Kazzaz, E. Bornstein, O. Lambotte, D. Altmann, B. R. Blazar, B. Rodriguez, L. Teixeira-Johnson, A. Landay, J. N. Martin, F. M. Hecht, L. J. Picker, M. M. Lederman, S. G. Deeks, D. C. Douek, Microbial translocation is a cause of systemic immune activation in chronic HIV infection. Nat Med 12, 1365–1371 (2006).

13. A. K. Steele, E. J. Lee, B. Vestal, D. Hecht, Z. Dong, E. Rapaport, J. Koeppe, T. B. Campbell, C. C. Wilson, Contribution of intestinal barrier damage, microbial translocation and HIV-1 infection status to an inflammaging signature. PLoS One 9, e97171 (2014).

14. A. S. Zevin, L. McKinnon, A. Burgener, N. R. Klatt, Microbial translocation and microbiome dysbiosis in HIV-associated immune activation. Curr Opin HIV AIDS 11, 182–190 (2016).

15. G. Ancona, E. Merlini, C. Tincati, A. Barassi, A. Calcagno, M. Augello, V. Bono, F. Bai, E. S. Cannizzo, A. d’Arminio Monforte, G. Marchetti, Long-Term Suppressive cART Is Not Sufficient to Restore Intestinal Permeability and Gut Microbiota Compositional Changes. Front Immunol 12, 639291 (2021).

16. J. D. Estes, L. D. Harris, N. R. Klatt, B. Tabb, S. Pittaluga, M. Paiardini, G. R. Barclay, J. Smedley, R. Pung, K. M. Oliveira, V. M. Hirsch, G. Silvestri, D. C. Douek, C. J. Miller, A. T. Haase, J. Lifson, J. M. Brenchley, Damaged intestinal epithelial integrity linked to microbial translocation in pathogenic simian immunodeficiency virus infections. PLoS Pathog 6, e1001052 (2010).

17. M. Somsouk, J. D. Estes, C. Deleage, R. M. Dunham, R. Albright, J. M. Inadomi, J. N. Martin, S. G. Deeks, J. M. McCune, P. W. Hunt, Gut epithelial barrier and systemic inflammation during chronic HIV infection. AIDS 29, 43–51 (2015).

18. A. J. Macpherson, K. D. McCoy, F. E. Johansen, P. Brandtzaeg, The immune geography of IgA induction and function. Mucosal Immunol 1, 11–22 (2008).

19. M. A. McGuckin, S. K. Linden, P. Sutton, T. H. Florin, Mucin dynamics and enteric pathogens. Nat Rev Microbiol 9, 265–278 (2011).

20. B. Corthesy, Multi-faceted functions of secretory IgA at mucosal surfaces. Front Immunol 4, 185 (2013).

21. B. Pietrzak, K. Tomela, A. Olejnik-Schmidt, A. Mackiewicz, M. Schmidt, Secretory IgA in Intestinal Mucosal Secretions as an Adaptive Barrier against Microbial Cells. Int J Mol Sci 21, (2020).

22. J. Kabbert, J. Benckert, T. Rollenske, T. C. A. Hitch, T. Clavel, V. Cerovic, H. Wardemann, O. Pabst, High microbiota reactivity of adult human intestinal IgA requires somatic mutations. J Exp Med 217, (2020).

23. R. Elgueta, M. J. Benson, V. C. de Vries, A. Wasiuk, Y. Guo, R. J. Noelle, Molecular mechanism and function of CD40/CD40L engagement in the immune system. Immunol Rev 229, 152–172 (2009).

24. A. Reboldi, T. I. Arnon, L. B. Rodda, A. Atakilit, D. Sheppard, J. G. Cyster, IgA production requires B cell interaction with subepithelial dendritic cells in Peyer’s patches. Science 352, aaf4822 (2016).

25. G. D. Victora, M. C. Nussenzweig, Germinal Centers. Annu Rev Immunol 40, 413–442 (2022).

26. J. P. Pereira, L. M. Kelly, J. G. Cyster, Finding the right niche: B-cell migration in the early phases of T-dependent antibody responses. Int Immunol 22, 413–419 (2010).

27. T. A. Schwickert, G. D. Victora, D. R. Fooksman, A. O. Kamphorst, M. R. Mugnier, A. D. Gitlin, M. L. Dustin, M. C. Nussenzweig, A dynamic T cell-limited checkpoint regulates affinity-dependent B cell entry into the germinal center. J Exp Med 208, 1243–1252 (2011).

28. J. G. Cyster, C. D. C. Allen, B Cell Responses: Cell Interaction Dynamics and Decisions. Cell 177, 524–540 (2019).

29. S. M. Dillon, E. J. Lee, C. V. Kotter, G. L. Austin, Z. Dong, D. K. Hecht, S. Gianella, B. Siewe, D. M. Smith, A. L. Landay, C. E. Robertson, D. N. Frank, C. C. Wilson, An altered intestinal mucosal microbiome in HIV-1 infection is associated with mucosal and systemic immune activation and endotoxemia. Mucosal Immunol 7, 983–994 (2014).

30. C. A. Lozupone, M. Li, T. B. Campbell, S. C. Flores, D. Linderman, M. J. Gebert, R. Knight, A. P. Fontenot, B. E. Palmer, Alterations in the gut microbiota associated with HIV-1 infection. Cell Host Microbe 14, 329–339 (2013).

31. I. Vujkovic-Cvijin, R. M. Dunham, S. Iwai, M. C. Maher, R. G. Albright, M. J. Broadhurst, R. D. Hernandez, M. M. Lederman, Y. Huang, M. Somsouk, S. G. Deeks, P. W. Hunt, S. V. Lynch, J. M. McCune, Dysbiosis of the gut microbiota is associated with HIV disease progression and tryptophan catabolism. Sci Transl Med 5, 193ra191 (2013).

32. D. M. Dinh, G. E. Volpe, C. Duffalo, S. Bhalchandra, A. K. Tai, A. V. Kane, C. A. Wanke, H. D. Ward, Intestinal microbiota, microbial translocation, and systemic inflammation in chronic HIV infection. J Infect Dis 211, 19–27 (2015).

33. J. F. Vazquez-Castellanos, S. Serrano-Villar, A. Latorre, A. Artacho, M. L. Ferrus, N. Madrid, A. Vallejo, T. Sainz, J. Martinez-Botas, S. Ferrando-Martinez, M. Vera, F. Dronda, M. Leal, J. Del Romero, S. Moreno, V. Estrada, M. J. Gosalbes, A. Moya, Altered metabolism of gut microbiota contributes to chronic immune activation in HIV-infected individuals. Mucosal Immunol 8, 760–772 (2015).

34. E. A. Mutlu, A. Keshavarzian, J. Losurdo, G. Swanson, B. Siewe, C. Forsyth, A. French, P. Demarais, Y. Sun, L. Koenig, S. Cox, P. Engen, P. Chakradeo, R. Abbasi, A. Gorenz, C. Burns, A. Landay, A compositional look at the human gastrointestinal microbiome and immune activation parameters in HIV infected subjects. PLoS Pathog 10, e1003829 (2014).

35. C. L. Monaco, D. B. Gootenberg, G. Zhao, S. A. Handley, M. S. Ghebremichael, E. S. Lim, A. Lankowski, M. T. Baldridge, C. B. Wilen, M. Flagg, J. M. Norman, B. C. Keller, J. M. Luevano, D. Wang, Y. Boum, J. N. Martin, P. W. Hunt, D. R. Bangsberg, M. J. Siedner, D. S. Kwon, H. W. Virgin, Altered Virome and Bacterial Microbiome in Human Immunodeficiency Virus-Associated Acquired Immunodeficiency Syndrome. Cell Host Microbe 19, 311–322 (2016).

36. W. Lu, Y. Feng, F. Jing, Y. Han, N. Lyu, F. Liu, J. Li, X. Song, J. Xie, Z. Qiu, T. Zhu, B. Routy, J. P. Routy, T. Li, B. Zhu, Association Between Gut Microbiota and CD4 Recovery in HIV-1 Infected Patients. Front Microbiol 9, 1451 (2018).

37. J. M. Brenchley, M. Paiardini, K. S. Knox, A. I. Asher, B. Cervasi, T. E. Asher, P. Scheinberg, D. A. Price, C. A. Hage, L. M. Kholi, A. Khoruts, I. Frank, J. Else, T. Schacker, G. Silvestri, D. C. Douek, Differential Th17 CD4 T-cell depletion in pathogenic and nonpathogenic lentiviral infections. Blood 112, 2826–2835 (2008).

38. V. Cecchinato, C. J. Trindade, A. Laurence, J. M. Heraud, J. M. Brenchley, M. G. Ferrari, L. Zaffiri, E. Tryniszewska, W. P. Tsai, M. Vaccari, R. W. Parks, D. Venzon, D. C. Douek, J. J. O’Shea, G. Franchini, Altered balance between Th17 and Th1 cells at mucosal sites predicts AIDS progression in simian immunodeficiency virus-infected macaques. Mucosal Immunol 1, 279–288 (2008).

39. S. M. Tugizov, R. Herrera, P. Chin-Hong, P. Veluppillai, D. Greenspan, J. Michael Berry, C. D. Pilcher, C. H. Shiboski, N. Jay, M. Rubin, A. Chein, J. M. Palefsky, HIV-associated disruption of mucosal epithelium facilitates paracellular penetration by human papillomavirus. Virology 446, 378–388 (2013).

40. P. Brandtzaeg, H. Kiyono, R. Pabst, M. W. Russell, Terminology: nomenclature of mucosa-associated lymphoid tissue. Mucosal Immunol 1, 31–37 (2008).

41. T. Stuart, A. Butler, P. Hoffman, C. Hafemeister, E. Papalexi, W. M. Mauck, 3rd, Y. Hao, M. Stoeckius, P. Smibert, R. Satija, Comprehensive Integration of Single-Cell Data. Cell 177, 1888–1902 e1821 (2019).

42. B. A. Heesters, K. van Megesen, I. Tomris, R. P. de Vries, G. Magri, H. Spits, Characterization of human FDCs reveals regulation of T cells and antigen presentation to B cells. J Exp Med 218, (2021).

43. O. J. Landsverk, O. Snir, R. B. Casado, L. Richter, J. E. Mold, P. Reu, R. Horneland, V. Paulsen, S. Yaqub, E. M. Aandahl, O. M. Oyen, H. S. Thorarensen, M. Salehpour, G. Possnert, J. Frisen, L. M. Sollid, E. S. Baekkevold, F. L. Jahnsen, Antibody-secreting plasma cells persist for decades in human intestine. J Exp Med 214, 309–317 (2017).

44. C. M. Buckner, S. Moir, J. Ho, W. Wang, J. G. Posada, L. Kardava, E. K. Funk, A. K. Nelson, Y. Li, T. W. Chun, A. S. Fauci, Characterization of plasmablasts in the blood of HIV-infected viremic individuals: evidence for nonspecific immune activation. J Virol 87, 5800–5811 (2013).

45. H. E. Mei, T. Yoshida, W. Sime, F. Hiepe, K. Thiele, R. A. Manz, A. Radbruch, T. Dorner, Blood-borne human plasma cells in steady state are derived from mucosal immune responses. Blood 113, 2461–2469 (2009).

46. S. R. Savage, Z. Shi, Y. Liao, B. Zhang, Graph Algorithms for Condensing and Consolidating Gene Set Analysis Results. Mol Cell Proteomics 18, S141–S152 (2019).

47. M. Perreau, A. L. Savoye, E. De Crignis, J. M. Corpataux, R. Cubas, E. K. Haddad, L. De Leval, C. Graziosi, G. Pantaleo, Follicular helper T cells serve as the major CD4 T cell compartment for HIV-1 infection, replication, and production. J Exp Med 210, 143–156 (2013).

48. E. Moysi, S. Pallikkuth, L. R. De Armas, L. E. Gonzalez, D. Ambrozak, V. George, D. Huddleston, R. Pahwa, R. A. Koup, C. Petrovas, S. Pahwa, Altered immune cell follicular dynamics in HIV infection following influenza vaccination. J Clin Invest 128, 3171–3185 (2018).

49. Z. Liu, W. G. Cumberland, L. E. Hultin, A. H. Kaplan, R. Detels, J. V. Giorgi, CD8+ T-lymphocyte activation in HIV-1 disease reflects an aspect of pathogenesis distinct from viral burden and immunodeficiency. J Acquir Immune Defic Syndr Hum Retrovirol 18, 332–340 (1998).

50. J. V. Giorgi, L. E. Hultin, J. A. McKeating, T. D. Johnson, B. Owens, L. P. Jacobson, R. Shih, J. Lewis, D. J. Wiley, J. P. Phair, S. M. Wolinsky, R. Detels, Shorter survival in advanced human immunodeficiency virus type 1 infection is more closely associated with T lymphocyte activation than with plasma virus burden or virus chemokine coreceptor usage. J Infect Dis 179, 859–870 (1999).

51. I. Sereti, M. Altfeld, Immune activation and HIV: an enduring relationship. Curr Opin HIV AIDS 11, 129–130 (2016).

52. L. A. van der Waaij, P. C. Limburg, G. Mesander, D. van der Waaij, In vivo IgA coating of anaerobic bacteria in human faeces. Gut 38, 348–354 (1996).

53. K. Suzuki, B. Meek, Y. Doi, M. Muramatsu, T. Chiba, T. Honjo, S. Fagarasan, Aberrant expansion of segmented filamentous bacteria in IgA-deficient gut. Proc Natl Acad Sci U S A 101, 1981–1986 (2004).

54. C. Gutzeit, G. Magri, A. Cerutti, Intestinal IgA production and its role in host-microbe interaction. Immunol Rev 260, 76–85 (2014).

55. A. Nakajima, A. Vogelzang, M. Maruya, M. Miyajima, M. Murata, A. Son, T. Kuwahara, T. Tsuruyama, S. Yamada, M. Matsuura, H. Nakase, D. A. Peterson, S. Fagarasan, K. Suzuki, IgA regulates the composition and metabolic function of gut microbiota by promoting symbiosis between bacteria. J Exp Med 215, 2019–2034 (2018).

56. O. Pabst, E. Slack, IgA and the intestinal microbiota: the importance of being specific. Mucosal Immunol 13, 12–21 (2020).

57. N. W. Palm, M. R. de Zoete, T. W. Cullen, N. A. Barry, J. Stefanowski, L. Hao, P. H. Degnan, J. Hu, I. Peter, W. Zhang, E. Ruggiero, J. H. Cho, A. L. Goodman, R. A. Flavell, Immunoglobulin A coating identifies colitogenic bacteria in inflammatory bowel disease. Cell 158, 1000–1010 (2014).

58. S. H. Duncan, A. Belenguer, G. Holtrop, A. M. Johnstone, H. J. Flint, G. E. Lobley, Reduced dietary intake of carbohydrates by obese subjects results in decreased concentrations of butyrate and butyrate-producing bacteria in feces. Appl Environ Microbiol 73, 1073–1078 (2007).

59. T. S. Ghosh, S. Rampelli, I. B. Jeffery, A. Santoro, M. Neto, M. Capri, E. Giampieri, A. Jennings, M. Candela, S. Turroni, E. G. Zoetendal, G. D. A. Hermes, C. Elodie, N. Meunier, C. M. Brugere, E. Pujos-Guillot, A. M. Berendsen, L. De Groot, E. J. M. Feskins, …, P. W. O’Toole, Mediterranean diet intervention alters the gut microbiome in older people reducing frailty and improving health status: the NU-AGE 1-year dietary intervention across five European countries. Gut 69, 1218–1228 (2020).

60. G. Jakobsdottir, J. Xu, G. Molin, S. Ahrne, M. Nyman, High-fat diet reduces the formation of butyrate, but increases succinate, inflammation, liver fat and cholesterol in rats, while dietary fibre counteracts these effects. PLoS One 8, e80476 (2013).

61. M. Noguera-Julian, M. Rocafort, Y. Guillen, J. Rivera, M. Casadella, P. Nowak, F. Hildebrand, G. Zeller, M. Parera, R. Bellido, C. Rodriguez, J. Carrillo, B. Mothe, J. Coll, I. Bravo, C. Estany, C. Herrero, J. Saz, G. Sirera, …, R. Paredes, Gut Microbiota Linked to Sexual Preference and HIV Infection. EBioMedicine 5, 135–146 (2016).

62. C. F. Kelley, C. S. Kraft, T. J. de Man, C. Duphare, H. W. Lee, J. Yang, K. A. Easley, G. K. Tharp, M. J. Mulligan, P. S. Sullivan, S. E. Bosinger, R. R. Amara, The rectal mucosa and condomless receptive anal intercourse in HIV-negative MSM: implications for HIV transmission and prevention. Mucosal Immunol 10, 996–1007 (2017).

63. A. J. S. Armstrong, M. Shaffer, N. M. Nusbacher, C. Griesmer, S. Fiorillo, J. M. Schneider, C. Preston Neff, S. X. Li, A. P. Fontenot, T. Campbell, B. E. Palmer, C. A. Lozupone, An exploration of Prevotella-rich microbiomes in HIV and men who have sex with men. Microbiome 6, 198 (2018).

64. A. D. Gitlin, Z. Shulman, M. C. Nussenzweig, Clonal selection in the germinal centre by regulated proliferation and hypermutation. Nature 509, 637–640 (2014).

65. A. D. Gitlin, L. von Boehmer, A. Gazumyan, Z. Shulman, T. Y. Oliveira, M. C. Nussenzweig, Independent Roles of Switching and Hypermutation in the Development and Persistence of B Lymphocyte Memory. Immunity 44, 769–781 (2016).

66. J. M. Tas, L. Mesin, G. Pasqual, S. Targ, J. T. Jacobsen, Y. M. Mano, C. S. Chen, J. C. Weill, C. A. Reynaud, E. P. Browne, M. Meyer-Hermann, G. D. Victora, Visualizing antibody affinity maturation in germinal centers. Science 351, 1048–1054 (2016).

67. J. Ersching, A. Efeyan, L. Mesin, J. T. Jacobsen, G. Pasqual, B. C. Grabiner, D. Dominguez-Sola, D. M. Sabatini, G. D. Victora, Germinal Center Selection and Affinity Maturation Require Dynamic Regulation of mTORC1 Kinase. Immunity 46, 1045–1058 e1046 (2017).

68. M. Pauthner, C. Havenar-Daughton, D. Sok, J. P. Nkolola, R. Bastidas, A. V. Boopathy, D. G. Carnathan, A. Chandrashekar, K. M. Cirelli, C. A. Cottrell, A. M. Eroshkin, J. Guenaga, K. Kaushik, D. W. Kulp, J. Liu, L. E. McCoy, A. L. Oom, G. Ozorowski, K. W. Post, …, D. R. Burton, Elicitation of Robust Tier 2 Neutralizing Antibody Responses in Nonhuman Primates by HIV Envelope Trimer Immunization Using Optimized Approaches. Immunity 46, 1073–1088 e1076 (2017).

69. B. F. Haynes, K. Wiehe, P. Borrow, K. O. Saunders, B. Korber, K. Wagh, A. J. McMichael, G. Kelsoe, B. H. Hahn, F. Alt, G. M. Shaw, Strategies for HIV-1 vaccines that induce broadly neutralizing antibodies. Nat Rev Immunol 23, 142–158 (2023).

70. A. M. Trama, M. A. Moody, S. M. Alam, F. H. Jaeger, B. Lockwood, R. Parks, K. E. Lloyd, C. Stolarchuk, R. Scearce, A. Foulger, D. J. Marshall, J. F. Whitesides, T. L. Jeffries, Jr., K. Wiehe, L. Morris, B. Lambson, K. Soderberg, K. K. Hwang, G. D. Tomaras, …, B. F. Haynes, HIV-1 envelope gp41 antibodies can originate from terminal ileum B cells that share cross-reactivity with commensal bacteria. Cell Host Microbe 16, 215–226 (2014).

71. C. Planchais, A. Kok, A. Kanyavuz, V. Lorin, T. Bruel, F. Guivel-Benhassine, T. Rollenske, J. Prigent, T. Hieu, T. Prazuck, L. Lefrou, H. Wardemann, O. Schwartz, J. D. Dimitrov, L. Hocqueloux, H. Mouquet, HIV-1 Envelope Recognition by Polyreactive and Cross-Reactive Intestinal B Cells. Cell Rep 27, 572–585 e577 (2019).

72. S. Clare, V. John, A. W. Walker, J. L. Hill, C. Abreu-Goodger, C. Hale, D. Goulding, T. D. Lawley, P. Mastroeni, G. Frankel, A. J. Enright, E. Vigorito, G. Dougan, Enhanced susceptibility to Citrobacter rodentium infection in microRNA-155-deficient mice. Infect Immun 81, 723–732 (2013).

73. W. T. Yewdell, R. M. Smolkin, K. T. Belcheva, A. Mendoza, A. J. Michaels, M. Cols, D. Angeletti, J. W. Yewdell, J. Chaudhuri, Temporal dynamics of persistent germinal centers and memory B cell differentiation following respiratory virus infection. Cell Rep 37, 109961 (2021).

74. T. Hagglof, M. Cipolla, M. Loewe, S. T. Chen, L. Mesin, H. Hartweger, M. A. ElTanbouly, A. Cho, A. Gazumyan, V. Ramos, L. Stamatatos, T. Y. Oliveira, M. C. Nussenzweig, C. Viant, Continuous germinal center invasion contributes to the diversity of the immune response. Cell 186, 147–161 e115 (2023).

75. A. P. Hendrickx, J. Top, J. R. Bayjanov, H. Kemperman, M. R. Rogers, F. L. Paganelli, M. J. Bonten, R. J. Willems, Antibiotic-Driven Dysbiosis Mediates Intraluminal Agglutination and Alternative Segregation of Enterococcus faecium from the Intestinal Epithelium. mBio 6, e01346–01315 (2015).

76. K. J. Levinson, M. De Jesus, N. J. Mantis, Rapid effects of a protective O-polysaccharide-specific monoclonal IgA on Vibrio cholerae agglutination, motility, and surface morphology. Infect Immun 83, 1674–1683 (2015).

77. W. Xu, P. A. Santini, J. S. Sullivan, B. He, M. Shan, S. C. Ball, W. B. Dyer, T. J. Ketas, A. Chadburn, L. Cohen-Gould, D. M. Knowles, A. Chiu, R. W. Sanders, K. Chen, A. Cerutti, HIV-1 evades virus-specific IgG2 and IgA responses by targeting systemic and intestinal B cells via long-range intercellular conduits. Nat Immunol 10, 1008–1017 (2009).

78. S. Kaw, S. Ananth, N. Tsopoulidis, K. Morath, B. M. Coban, R. Hohenberger, O. C. Bulut, F. Klein, B. Stolp, O. T. Fackler, HIV-1 infection of CD4 T cells impairs antigen-specific B cell function. EMBO J 39, e105594 (2020).

79. P. Eldin, S. Peron, A. Galashevskaya, N. Denis-Lagache, M. Cogne, G. Slupphaug, L. Briant, Impact of HIV-1 Vpr manipulation of the DNA repair enzyme UNG2 on B lymphocyte class switch recombination. J Transl Med 18, 310 (2020).

80. L. Manganaro, E. de Castro, A. M. Maestre, K. Olivieri, A. Garcia-Sastre, A. Fernandez-Sesma, V. Simon, HIV Vpu Interferes with NF-kappaB Activity but Not with Interferon Regulatory Factor 3. J Virol 89, 9781–9790 (2015).

81. M. Lindqvist, J. van Lunzen, D. Z. Soghoian, B. D. Kuhl, S. Ranasinghe, G. Kranias, M. D. Flanders, S. Cutler, N. Yudanin, M. I. Muller, I. Davis, D. Farber, P. Hartjen, F. Haag, G. Alter, J. Schulze zur Wiesch, H. Streeck, Expansion of HIV-specific T follicular helper cells in chronic HIV infection. J Clin Invest 122, 3271–3280 (2012).

82. R. A. Cubas, J. C. Mudd, A. L. Savoye, M. Perreau, J. van Grevenynghe, T. Metcalf, E. Connick, A. Meditz, G. J. Freeman, G. Abesada-Terk, Jr., J. M. Jacobson, A. D. Brooks, S. Crotty, J. D. Estes, G. Pantaleo, M. M. Lederman, E. K. Haddad, Inadequate T follicular cell help impairs B cell immunity during HIV infection. Nat Med 19, 494–499 (2013).

83. M. C. Levesque, M. A. Moody, K. K. Hwang, D. J. Marshall, J. F. Whitesides, J. D. Amos, T. C. Gurley, S. Allgood, B. B. Haynes, N. A. Vandergrift, S. Plonk, D. C. Parker, M. S. Cohen, G. D. Tomaras, P. A. Goepfert, G. M. Shaw, J. E. Schmitz, J. J. Eron, N. J. Shaheen, …, B. F. Haynes, Polyclonal B cell differentiation and loss of gastrointestinal tract germinal centers in the earliest stages of HIV-1 infection. PLoS Med 6, e1000107 (2009).

84. J. D. Estes, Pathobiology of HIV/SIV-associated changes in secondary lymphoid tissues. Immunol Rev 254, 65–77 (2013).

85. M. A. Piris, C. Rivas, M. Morente, C. Rubio, C. Martin, H. Olivia, Persistent and generalized lymphadenopathy: a lesion of follicular dendritic cells? An immunohistologic and ultrastructural study. Am J Clin Pathol 87, 716–724 (1987).

86. K. Tenner-Racz, Human immunodeficiency virus associated changes in germinal centers of lymph nodes and relevance to impaired B-cell function. Lymphology 21, 36–43 (1988).

87. J. Schmitz, J. van Lunzen, K. Tenner-Racz, G. Grossschupff, P. Racz, H. Schmitz, M. Dietrich, F. T. Hufert, Follicular dendritic cells retain HIV-1 particles on their plasma membrane, but are not productively infected in asymptomatic patients with follicular hyperplasia. J Immunol 153, 1352–1359 (1994).

88. O. Devergne, M. Peuchmaur, M. C. Crevon, J. A. Trapani, M. C. Maillot, P. Galanaud, D. Emilie, Activation of cytotoxic cells in hyperplastic lymph nodes from HIV-infected patients. AIDS 5, 1071–1079 (1991).

89. S. W. Craig, J. J. Cebra, Peyer’s patches: an enriched source of precursors for IgA-producing immunocytes in the rabbit. J Exp Med 134, 188–200 (1971).

90. P. Brandtzaeg, Presence of J chain in human immunocytes containing various immunoglobulin classes. Nature 252, 418–420 (1974).

91. A. Cerutti, K. Chen, A. Chorny, Immunoglobulin responses at the mucosal interface. Annu Rev Immunol 29, 273–293 (2011).

92. N. Y. Lycke, M. Bemark, The regulation of gut mucosal IgA B-cell responses: recent developments. Mucosal Immunol 10, 1361–1374 (2017).

93. C. M. Snapper, W. E. Paul, Interferon-gamma and B cell stimulatory factor-1 reciprocally regulate Ig isotype production. Science 236, 944–947 (1987).

94. T. D. Langford, M. P. Housley, M. Boes, J. Chen, M. F. Kagnoff, F. D. Gillin, L. Eckmann, Central importance of immunoglobulin A in host defense against Giardia spp. Infect Immun 70, 11–18 (2002).

95. K. Endt, B. Stecher, S. Chaffron, E. Slack, N. Tchitchek, A. Benecke, L. Van Maele, J. C. Sirard, A. J. Mueller, M. Heikenwalder, A. J. Macpherson, R. Strugnell, C. von Mering, W. D. Hardt, The microbiota mediates pathogen clearance from the gut lumen after non-typhoidal Salmonella diarrhea. PLoS Pathog 6, e1001097 (2010).

96. D. O. Matson, M. L. O’Ryan, I. Herrera, L. K. Pickering, M. K. Estes, Fecal antibody responses to symptomatic and asymptomatic rotavirus infections. J Infect Dis 167, 577–583 (1993).

97. N. Feng, J. W. Burns, L. Bracy, H. B. Greenberg, Comparison of mucosal and systemic humoral immune responses and subsequent protection in mice orally inoculated with a homologous or a heterologous rotavirus. J Virol 68, 7766–7773 (1994).

98. S. E. Blutt, M. E. Conner, The gastrointestinal frontier: IgA and viruses. Front Immunol 4, 402 (2013).

99. J. Zaunders, M. Danta, M. Bailey, G. Mak, K. Marks, N. Seddiki, Y. Xu, D. J. Templeton, D. A. Cooper, M. A. Boyd, A. D. Kelleher, K. K. Koelsch, CD4(+) T Follicular Helper and IgA(+) B Cell Numbers in Gut Biopsies from HIV-Infected Subjects on Antiretroviral Therapy Are Similar to HIV-Uninfected Individuals. Front Immunol 7, 438 (2016).

100. C. Planchais, L. Hocqueloux, C. Ibanez, S. Gallien, C. Copie, M. Surenaud, A. Kok, V. Lorin, M. Fusaro, M. H. Delfau-Larue, L. Lefrou, T. Prazuck, M. Levy, N. Seddiki, J. D. Lelievre, H. Mouquet, Y. Levy, S. Hue, Early Antiretroviral Therapy Preserves Functional Follicular Helper T and HIV-Specific B Cells in the Gut Mucosa of HIV-1-Infected Individuals. J Immunol 200, 3519–3529 (2018).

101. A. Kok, L. Hocqueloux, H. Hocini, M. Carriere, L. Lefrou, A. Guguin, P. Tisserand, H. Bonnabau, V. Avettand-Fenoel, T. Prazuck, S. Katsahian, P. Gaulard, R. Thiebaut, Y. Levy, S. Hue, Early initiation of combined antiretroviral therapy preserves immune function in the gut of HIV-infected patients. Mucosal Immunol 8, 127–140 (2015).

102. D. J. Morrison, T. Preston, Formation of short chain fatty acids by the gut microbiota and their impact on human metabolism. Gut Microbes 7, 189–200 (2016).

103. V. Singh, G. Lee, H. Son, H. Koh, E. S. Kim, T. Unno, J. H. Shin, Butyrate producers, “The Sentinel of Gut”: Their intestinal significance with and beyond butyrate, and prospective use as microbial therapeutics. Front Microbiol 13, 1103836 (2022).

104. R. R. Fagundes, S. C. Belt, B. M. Bakker, G. Dijkstra, H. J. M. Harmsen, K. N. Faber, Beyond butyrate: microbial fiber metabolism supporting colonic epithelial homeostasis. Trends Microbiol, (2023).

105. M. M. Seldin, Y. Meng, H. Qi, W. Zhu, Z. Wang, S. L. Hazen, A. J. Lusis, D. M. Shih, Trimethylamine N-Oxide Promotes Vascular Inflammation Through Signaling of Mitogen-Activated Protein Kinase and Nuclear Factor-kappaB. J Am Heart Assoc 5, (2016).

106. B. C. Fu, M. A. J. Hullar, T. W. Randolph, A. A. Franke, K. R. Monroe, I. Cheng, L. R. Wilkens, J. A. Shepherd, M. M. Madeleine, L. Le Marchand, U. Lim, J. W. Lampe, Associations of plasma trimethylamine N-oxide, choline, carnitine, and betaine with inflammatory and cardiometabolic risk biomarkers and the fecal microbiome in the Multiethnic Cohort Adiposity Phenotype Study. Am J Clin Nutr 111, 1226–1234 (2020).

107. X. Liu, Y. Shao, J. Tu, J. Sun, L. Li, J. Tao, J. Chen, Trimethylamine-N-oxide-stimulated hepatocyte-derived exosomes promote inflammation and endothelial dysfunction through nuclear factor-kappa B signaling. Ann Transl Med 9, 1670 (2021).

108. S. F. Lima, L. Gogokhia, M. Viladomiu, L. Chou, G. Putzel, W. B. Jin, S. Pires, C. J. Guo, Y. Gerardin, C. V. Crawford, V. Jacob, E. Scherl, S. E. Brown, J. Hambor, R. S. Longman, Transferable Immunoglobulin A-Coated Odoribacter splanchnicus in Responders to Fecal Microbiota Transplantation for Ulcerative Colitis Limits Colonic Inflammation. Gastroenterology 162, 166–178 (2022).

109. M. Osawa, O. Handa, S. Fukushima, H. Matsumoto, E. Umegaki, R. Inoue, Y. Naito, A. Shiotani, Reduced abundance of butyric acid-producing bacteria in the ileal mucosa-associated microbiota of ulcerative colitis patients. J Clin Biochem Nutr 73, 77–83 (2023).

110. T. Wang, P. R. Sternes, X. K. Guo, H. Zhao, C. Xu, H. Xu, Autoimmune diseases exhibit shared alterations in the gut microbiota. Rheumatology (Oxford), (2023).

111. F. Asnicar, S. E. Berry, A. M. Valdes, L. H. Nguyen, G. Piccinno, D. A. Drew, E. Leeming, R. Gibson, C. Le Roy, H. A. Khatib, L. Francis, M. Mazidi, O. Mompeo, M. Valles-Colomer, A. Tett, F. Beghini, L. Dubois, D. Bazzani, A. M. Thomas, …, N. Segata, Microbiome connections with host metabolism and habitual diet from 1,098 deeply phenotyped individuals. Nat Med 27, 321–332 (2021).

112. R. Gacesa, A. Kurilshikov, A. Vich Vila, T. Sinha, M. A. Y. Klaassen, L. A. Bolte, S. Andreu-Sanchez, L. Chen, V. Collij, S. Hu, J. A. M. Dekens, V. C. Lenters, J. R. Bjork, J. C. Swarte, M. A. Swertz, B. H. Jansen, J. Gelderloos-Arends, S. Jankipersadsing, M. Hofker, …, R. K. Weersma, Environmental factors shaping the gut microbiome in a Dutch population. Nature 604, 732–739 (2022).

113. I. H. McHardy, X. Li, M. Tong, P. Ruegger, J. Jacobs, J. Borneman, P. Anton, J. Braun, HIV Infection is associated with compositional and functional shifts in the rectal mucosal microbiota. Microbiome 1, 26 (2013).

114. I. Vujkovic-Cvijin, O. Sortino, E. Verheij, J. Sklar, F. W. Wit, N. A. Kootstra, B. Sellers, J. M. Brenchley, J. Ananworanich, M. S. V. Loeff, Y. Belkaid, P. Reiss, I. Sereti, HIV-associated gut dysbiosis is independent of sexual practice and correlates with noncommunicable diseases. Nat Commun 11, 2448 (2020).

115. I. Vujkovic-Cvijin, H. C. Welles, C. W. Y. Ha, L. Huq, S. Mistry, J. M. Brenchley, G. Trinchieri, S. Devkota, Y. Belkaid, The systemic anti-microbiota IgG repertoire can identify gut bacteria that translocate across gut barrier surfaces. Sci Transl Med 14, eabl3927 (2022).

116. T. Senda, P. Dogra, T. Granot, K. Furuhashi, M. E. Snyder, D. J. Carpenter, P. A. Szabo, P. Thapa, M. Miron, D. L. Farber, Microanatomical dissection of human intestinal T-cell immunity reveals site-specific changes in gut-associated lymphoid tissues over life. Mucosal Immunol 12, 378–389 (2019).

117. J. Spencer, J. H. Y. Siu, L. Montorsi, Human intestinal lymphoid tissue in time and space. Mucosal Immunol 12, 296–298 (2019).

118. T. M. Fenton, P. B. Jørgensen, K. Niss, S. J. S. Rubin, U. M. Mörbe, L. B. Riis, C. Da Silva, A. Plumb, J. Vandamme, H. L. Jakobsen, S. Brunak, A. Habtezion, O. H. Nielsen, B. Johansson-Lindbom, W. W. Agace, Immune Profiling of Human Gut-Associated Lymphoid Tissue Identifies a Role for Isolated Lymphoid Follicles in Priming of Region-Specific Immunity. Immunity 52, 557–570.e556 (2020).

119. G. Magri, L. Comerma, M. Pybus, J. Sintes, D. Llige, D. Segura-Garzon, S. Bascones, A. Yeste, E. K. Grasset, C. Gutzeit, M. Uzzan, M. Ramanujam, M. C. van Zelm, R. Albero-Gonzalez, I. Vazquez, M. Iglesias, S. Serrano, L. Marquez, E. Mercade, S. Mehandru, A. Cerutti, Human Secretory IgM Emerges from Plasma Cells Clonally Related to Gut Memory B Cells and Targets Highly Diverse Commensals. Immunity 47, 118–134 e118 (2017).

120. M. Uzzan, M. Tokuyama, A. K. Rosenstein, C. Tomescu, I. N. SahBandar, H. M. Ko, L. Leyre, A. Chokola, E. Kaplan-Lewis, G. Rodriguez, A. Seki, M. J. Corley, J. Aberg, A. La Porte, E. Y. Park, H. Ueno, I. Oikonomou, I. Doron, I. D. Iliev, …, S. Mehandru, Anti-α4β7 therapy targets lymphoid aggregates in the gastrointestinal tract of HIV-1-infected individuals. Sci Transl Med 10, (2018).

121. P. Canales-Herrerias, M. Uzzan, A. Seki, R. S. Czepielewski, B. Verstockt, A. Livanos, F. Raso, A. Dunn, D. Dai, A. Wang, Z. Al-Taie, J. Martin, H. M. Ko, M. Tokuyama, M. Tankelevich, H. Meringer, F. Cossarini, D. Jha, A. Krek, J. D. Paulsen, M. Z. Nakadar, J. Wong, E. C. Erlich, E. J. Onufer, B. A. Helmink, K. Sharma, A. Rosenstein, G. Chung, T. Dawson, J. Juarez, V. Yajnik, A. Cerutti, J. Faith, M. Suarez-Farinas, C. Argmann, F. Petralia, G. J. Randolph, A. D. Polydorides, A. Reboldi, J. F. Colombel, S. Mehandru, Gut-associated lymphoid tissue attrition associates with response to anti-alpha4beta7 therapy in ulcerative colitis. bioRxiv, (2023).

122. C. W. Law, Y. Chen, W. Shi, G. K. Smyth, voom: Precision weights unlock linear model analysis tools for RNA-seq read counts. Genome Biol 15, R29 (2014).

123. C. W. Law, M. Alhamdoosh, S. Su, X. Dong, L. Tian, G. K. Smyth, M. E. Ritchie, RNA-seq analysis is easy as 1-2-3 with limma, Glimma and edgeR. F1000Res 5, (2016).

124. M. E. Ritchie, B. Phipson, D. Wu, Y. Hu, C. W. Law, W. Shi, G. K. Smyth, limma powers differential expression analyses for RNA-sequencing and microarray studies. Nucleic Acids Res 43, e47 (2015).

125. B. Phipson, S. Lee, I. J. Majewski, W. S. Alexander, G. K. Smyth, Robust Hyperparameter Estimation Protects against Hypervariable Genes and Improves Power to Detect Differential Expression. Ann Appl Stat 10, 946–963 (2016).

126. G. K. Smyth, Linear models and empirical bayes methods for assessing differential expression in microarray experiments. Stat Appl Genet Mol Biol 3, Article3 (2004).

127. 10xgenomics.

128. Y. Hao, S. Hao, E. Andersen-Nissen, W. M. Mauck, 3rd, S. Zheng, A. Butler, M. J. Lee, A. J. Wilk, C. Darby, M. Zager, P. Hoffman, M. Stoeckius, E. Papalexi, E. P. Mimitou, J. Jain, A. Srivastava, T. Stuart, L. M. Fleming, B. Yeung, …, R. Satija, Integrated analysis of multimodal single-cell data. Cell 184, 3573–3587 e3529 (2021).

129. J. S. Weinstein, K. Lezon-Geyda, Y. Maksimova, S. Craft, Y. Zhang, M. Su, V. P. Schulz, J. Craft, P. G. Gallagher, Global transcriptome analysis and enhancer landscape of human primary T follicular helper and T effector lymphocytes. Blood 124, 3719–3729 (2014).

130. P. A. Szabo, H. M. Levitin, M. Miron, M. E. Snyder, T. Senda, J. Yuan, Y. L. Cheng, E. C. Bush, P. Dogra, P. Thapa, D. L. Farber, P. A. Sims, Single-cell transcriptomics of human T cells reveals tissue and activation signatures in health and disease. Nat Commun 10, 4706 (2019).

131. A. B. Holmes, C. Corinaldesi, Q. Shen, R. Kumar, N. Compagno, Z. Wang, M. Nitzan, E. Grunstein, L. Pasqualucci, R. Dalla-Favera, K. Basso, Single-cell analysis of germinal-center B cells informs on lymphoma cell of origin and outcome. J Exp Med 217, (2020).

132. S. W. Kazer, T. P. Aicher, D. M. Muema, S. L. Carroll, J. Ordovas-Montanes, V. N. Miao, A. A. Tu, C. G. K. Ziegler, S. K. Nyquist, E. B. Wong, N. Ismail, M. Dong, A. Moodley, B. Berger, J. C. Love, K. L. Dong, A. Leslie, Z. M. Ndhlovu, T. Ndung’u, …, A. K. Shalek, Integrated single-cell analysis of multicellular immune dynamics during hyperacute HIV-1 infection. Nat Med 26, 511–518 (2020).

133. H. W. King, N. Orban, J. C. Riches, A. J. Clear, G. Warnes, S. A. Teichmann, L. K. James, Single-cell analysis of human B cell maturation predicts how antibody class switching shapes selection dynamics. Sci Immunol 6, (2021).

134. M. Uzzan, J. C. Martin, L. Mesin, A. E. Livanos, T. Castro-Dopico, R. Huang, F. Petralia, G. Magri, S. Kumar, Q. Zhao, A. K. Rosenstein, M. Tokuyama, K. Sharma, R. Ungaro, R. Kosoy, D. Jha, J. Fischer, H. Singh, M. E. Keir, …, S. Mehandru, Ulcerative colitis is characterized by a plasmablast-skewed humoral response associated with disease activity. Nat Med 28, 766–779 (2022).

135. J. Santos, M. Embrechts, On the Use of the Adjusted Rand Index as a Metric for Evaluating Supervised Classification. (2009), vol. 2009, pp. 175–184.

136. A. Subramanian, P. Tamayo, V. K. Mootha, S. Mukherjee, B. L. Ebert, M. A. Gillette, A. Paulovich, S. L. Pomeroy, T. R. Golub, E. S. Lander, J. P. Mesirov, Gene set enrichment analysis: a knowledge-based approach for interpreting genome-wide expression profiles. Proc Natl Acad Sci U S A 102, 15545–15550 (2005).

137. D. Croft, G. O’Kelly, G. Wu, R. Haw, M. Gillespie, L. Matthews, M. Caudy, P. Garapati, G. Gopinath, B. Jassal, S. Jupe, I. Kalatskaya, S. Mahajan, B. May, N. Ndegwa, E. Schmidt, V. Shamovsky, C. Yung, E. Birney, …, L. Stein, Reactome: a database of reactions, pathways and biological processes. Nucleic Acids Res 39, D691–697 (2011).

138. W. Shi, Y. Liao, S. N. Willis, N. Taubenheim, M. Inouye, D. M. Tarlinton, G. K. Smyth, P. D. Hodgkin, S. L. Nutt, L. M. Corcoran, Transcriptional profiling of mouse B cell terminal differentiation defines a signature for antibody-secreting plasma cells. Nat Immunol 16, 663–673 (2015).

139. O. Snir, C. Kanduri, K. E. A. Lundin, G. K. Sandve, L. M. Sollid, Transcriptional profiling of human intestinal plasma cells reveals effector functions beyond antibody production. United European Gastroenterol J 7, 1399–1407 (2019).

140. A. Reyes, M. Wu, N. P. McNulty, F. L. Rohwer, J. I. Gordon, Gnotobiotic mouse model of phage-bacterial host dynamics in the human gut. Proc Natl Acad Sci U S A 110, 20236–20241 (2013).

141. D. T. Truong, E. A. Franzosa, T. L. Tickle, M. Scholz, G. Weingart, E. Pasolli, A. Tett, C. Huttenhower, N. Segata, MetaPhlAn2 for enhanced metagenomic taxonomic profiling. Nat Methods 12, 902–903 (2015).

142. N. Segata, J. Izard, L. Waldron, D. Gevers, L. Miropolsky, W. S. Garrett, C. Huttenhower, Metagenomic biomarker discovery and explanation. Genome Biol 12, R60 (2011).

143. P. J. McMurdie, S. Holmes, phyloseq: an R package for reproducible interactive analysis and graphics of microbiome census data. PLoS One 8, e61217 (2013).

144. E. J. Contijoch, G. J. Britton, C. Yang, I. Mogno, Z. Li, R. Ng, S. R. Llewellyn, S. Hira, C. Johnson, K. M. Rabinowitz, R. Barkan, I. Dotan, R. P. Hirten, S. C. Fu, Y. Luo, N. Yang, T. Luong, P. R. Labrias, S. Lira, …, J. J. Faith, Gut microbiota density influences host physiology and is shaped by host and microbial factors. Elife 8, (2019).

145. J. J. Faith, J. L. Guruge, M. Charbonneau, S. Subramanian, H. Seedorf, A. L. Goodman, J. C. Clemente, R. Knight, A. C. Heath, R. L. Leibel, M. Rosenbaum, J. I. Gordon, The long-term stability of the human gut microbiota. Science 341, 1237439 (2013).

