## Supplementary Figures for "Gastrointestinal germinal center B cell depletion and reduction in IgA^+^ plasma cells in HIV-1 infection"

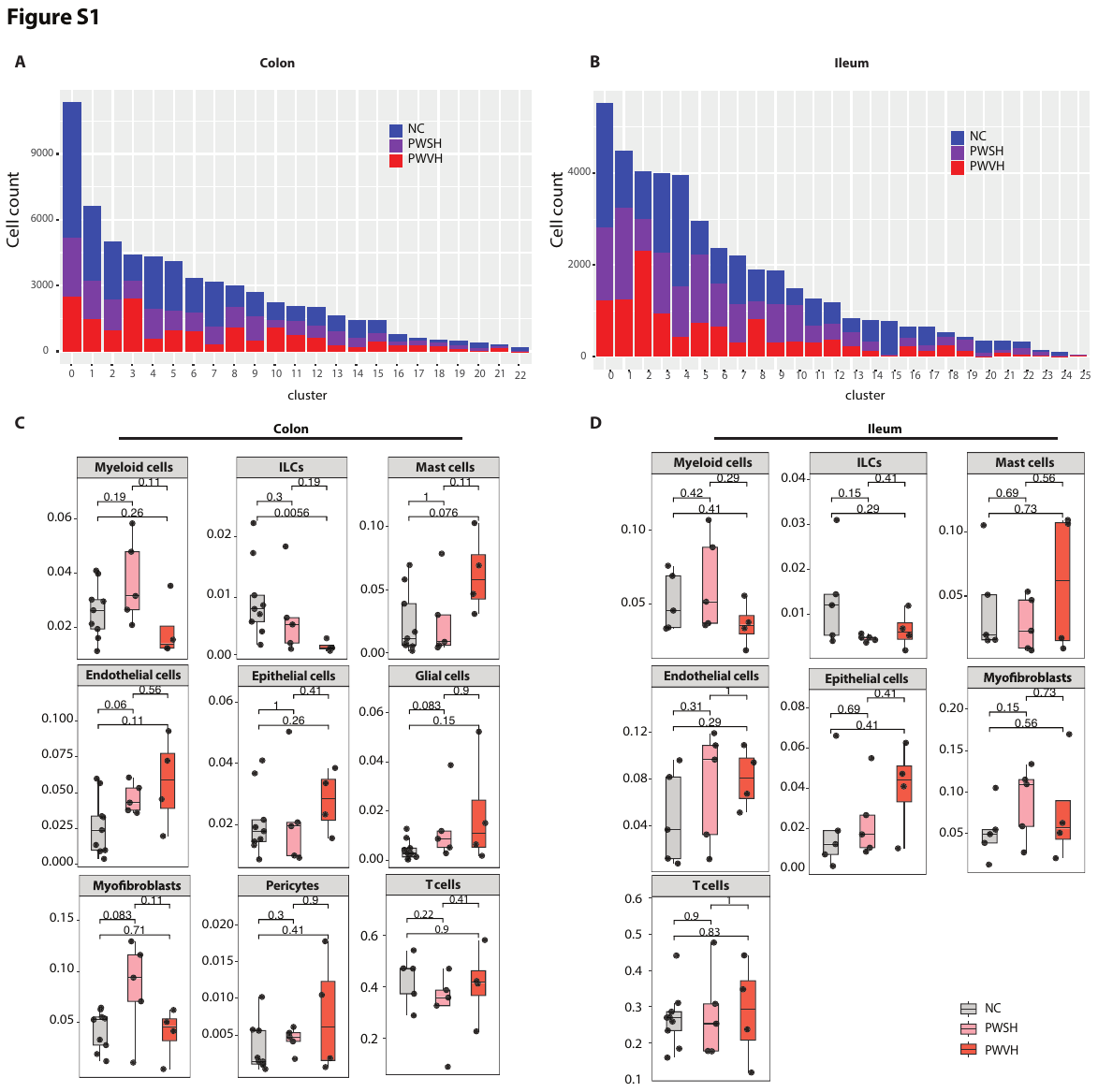

**Figure S1: scRNA-seq analyses of colonic and ileal tissue. (A)** Histogram showing cell count per colonic cell cluster. **(B)** Histogram showing cell count per ileal cell cluster. **(C)** Cell type distribution across the three groups of study participants in colonic samples (HIV negative controls (NC) n=9; people with suppressed HIV infection (PWSH) n=5; people with viremic HIV infection (PWVH) n=4). **(D)** Cell type distribution across the three groups of study participants in ileum-derived samples (NC, n=5; PWSH, n=5; PWVH, n=4). In panels C and D comparisons were performed with Wilcoxon test, p-values as indicated.

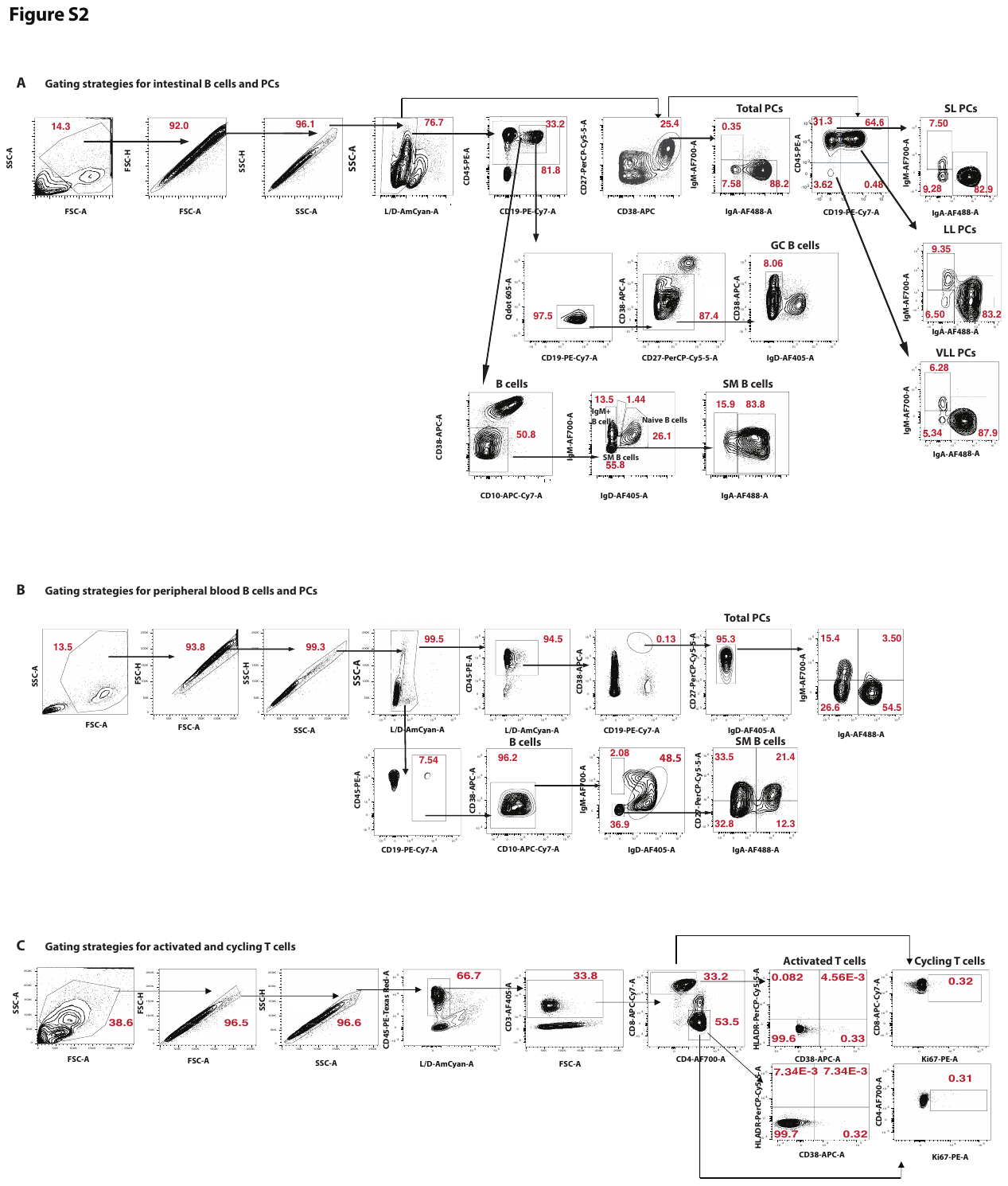

**Figure S2: Gating Strategy for flow cytometry data.** **(A)** Gating strategy used for intestinal B cells and plasma cells (PCs) showing PCs as live, CD27^+^CD38^hi^cells; Short-lived (SL) PCs as live, CD27^+^CD38^hi^CD45^+^CD19^+^cells, Long-lived (LL) PCs as live, D27^+^CD38^hi^CD45^+^CD19^-^cells, and Very Long-Lived (VLL) PCs as live, CD27^+^CD38^hi^CD45^-^CD19^-^cells. Total B cells are defined as live CD45^+^CD19^+^CD10^-^CD38^-^ and Germinal Center (GC) B cells are defined as CD45^+^CD3^-^CD19^+^CD38^int^IgD^-^ cells. Naïve B cells are defined as live CD45^+^CD19^+^CD10^-^CD38^-^IgM^+^IgD^+^ cells. Memory IgM expressing B Cells are defined as live CD45^+^CD19^+^CD10^-^CD38^-^IgM^+^ cells, Switched Memory (SM) B cells are defined as live CD45^+^CD19^+^CD10^-^CD38^-^IgM^-^IgD^-^cells. SM B cells are further classified into IgA^+^(live CD45^+^CD19^+^CD10^-^CD38^-^IgM^-^IgD^-^IgA^+^) and IgA^-^ (live CD45^+^CD19^+^CD10^-^CD38^-^IgM^-^IgD^-^ IgA) **(B)** Gating strategy used for peripheral blood (PB)-derived B cells and PCs showing PCs as live CD45^+^CD27^+^CD38^hi^IgD^-^D27^+^ cells and total B cells as live CD45^+^CD19^+^CD10^-^CD38^-^cells. Naïve B cells are defined as live CD45^+^CD19^+^CD10^-^CD38^-^IgM^+^IgD^+^ cells. Memory IgM expressing B cells are defined as live CD45^+^CD19^+^CD10^-^CD38^-^IgM^+^ cells, SM B Cells are defined as live CD45^+^CD19^+^CD10^-^CD38^-^IgM^-^IgD^-^cells. Flow plots pertaining to PB PCs are also shown in Fig. 3G. **(C)** Gating strategy used for activated and cycling T cells. CD4^+^ T cells are defined as live CD45^+^CD3^+^CD4^+^ cells. Similarly, CD8^+^ T cells are defined as liveCD45^+^CD3^+^CD8+cells. Activated CD4^+^ or CD8^+^ T cells are defined as CD38^+^ only, HLADR^+^ only or CD38^+^HLADR^+^ cells. Cycling T cells are defined as Ki67^+^ CD4^+^ or CD8^+^ T cells.

**Figure S3: Expression of CD4, BCL6 and CD23 in all NC samples.** Immunohistochemistry (IHC) images of intestinal lymphoid aggregates from all HIV Negative Controls showing the expression of CD4 (left panels), BCL6 (middle panels) and CD23 (right panels). Scale bar represents 100μm. Images from subject NV157 are depicted as representative of NC in Fig. 2c.

**Figure S4:** **Expression of CD4, BCL6 and CD23 in all PWVH samples.** Immunohistochemistry (IHC) images of intestinal lymphoid aggregates from all PWVH showing the expression of CD4 (left panels), BCL6 (middle panels) and CD23 (right panels). Scale bar represents 100μm. Images from subject HIV064 are depicted as representative of PWVH in Fig. 2c.

**Figure S5: Expression of CD4, BCL6 and CD23 in all PWSH samples.** Immunohistochemistry (IHC) images of intestinal lymphoid aggregates from all PWSH showing the expression of CD4 (left panels), BCL6 (middle panels) and CD23 (right panels). Scale bar represents 100μm. Images from subject HIV018 are depicted as representative of PWSH in Fig. 2c.

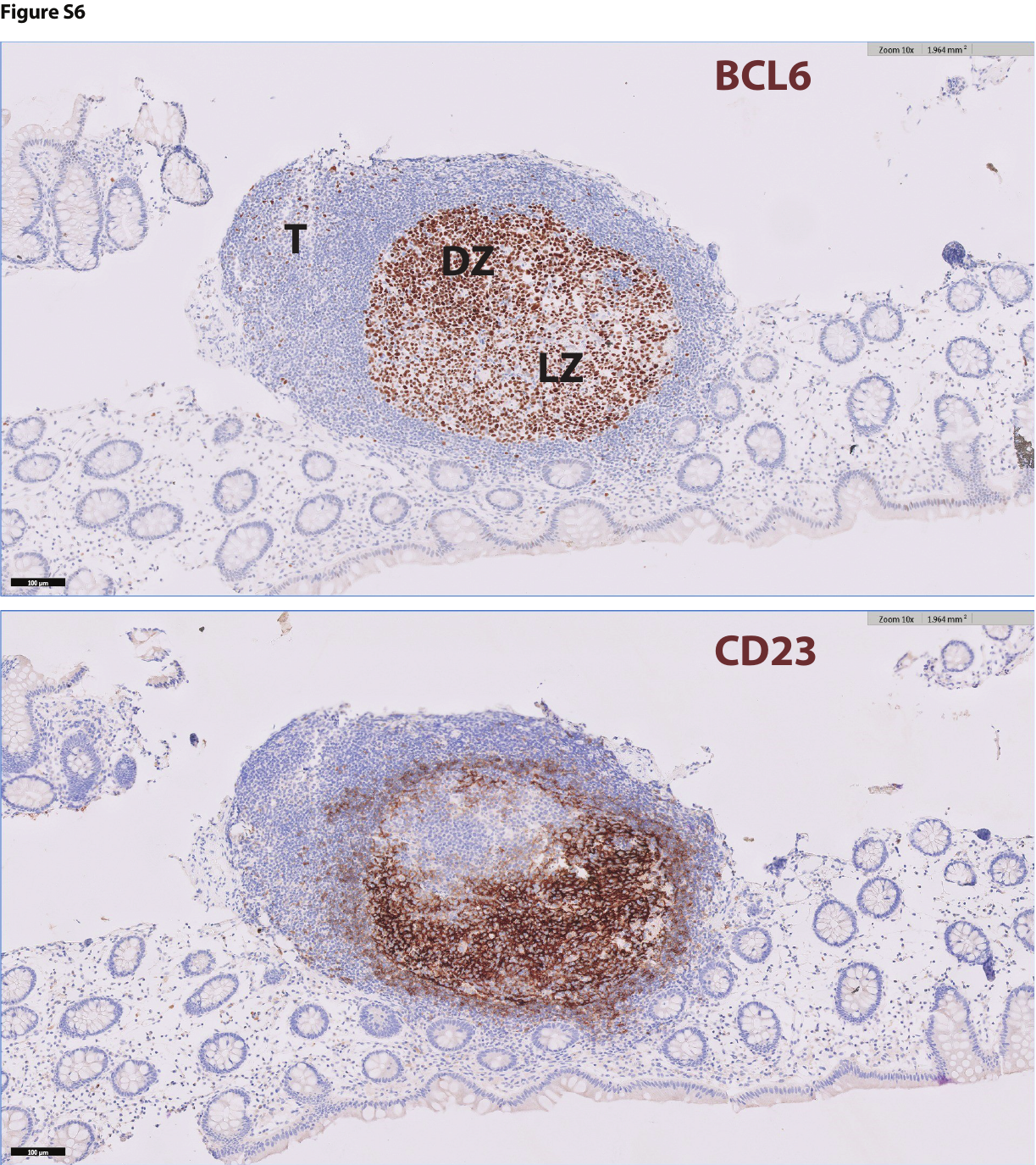

**Figure S6: Expression of CD23 in the putative light zones of the GCs.** Representative Immunohistochemistry (IHC) images of intestinal lymphoid aggregates from a NC showing the expression of BCL6 (top panel) and CD23 (bottom panel). LZ= light zone, DZ= dark zone, T= T cell zone. Scale bar represents 100μm.

**Figure S7: Expression of BCL6 correlates with CD23 in PWVH.** Representative IHC images from a PWVH (HIV066) showing 4 different lymphoid aggregates, comparing the expression of BCL6 and CD23. Yellow arrows indicate enhanced expression of BCL6 in a lymphoid follicle that demonstrated CD23 staining.

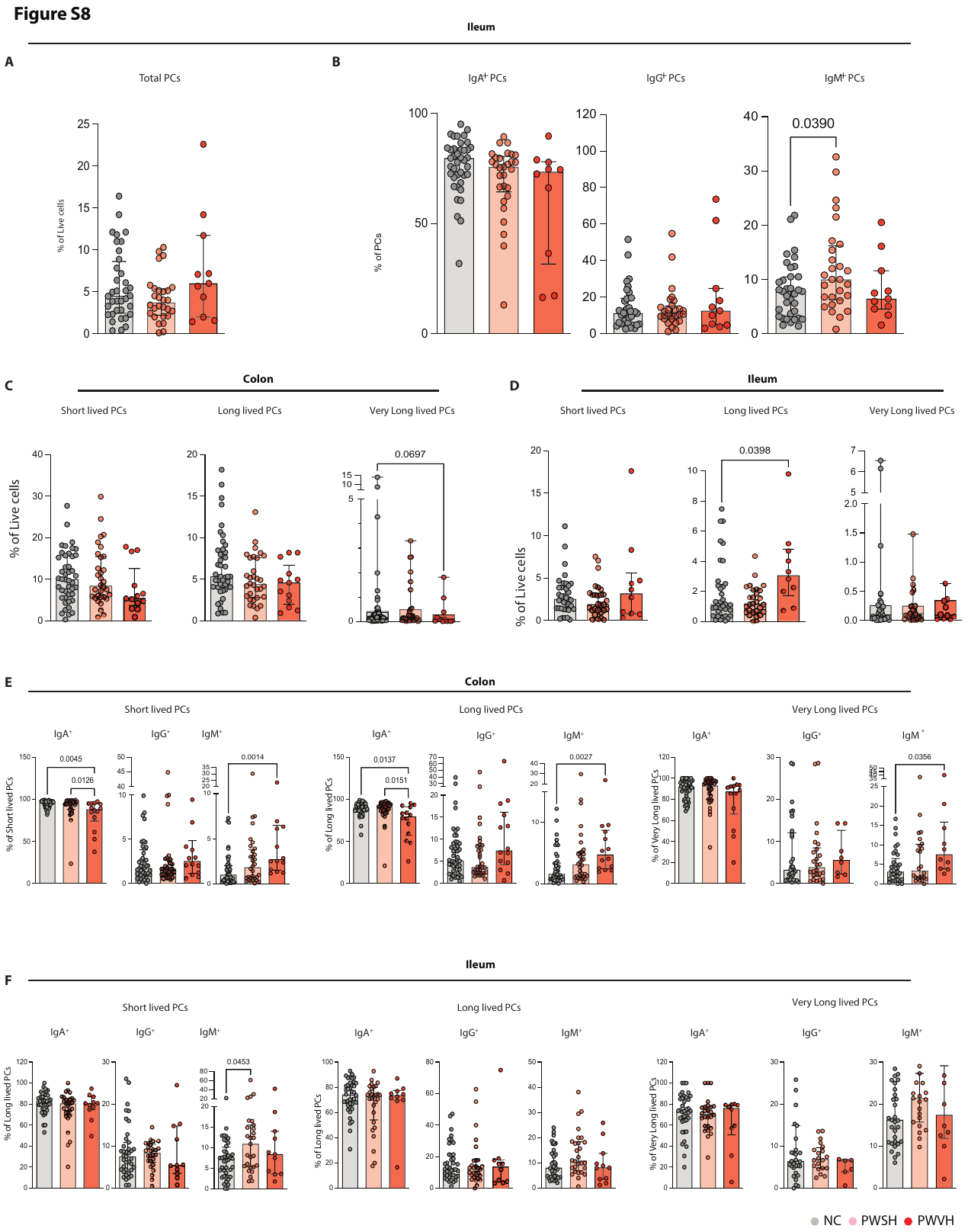

**Figure S8: Flow cytometric analyses demonstrate reduced PCs and isotype changes across different subtypes of PCs. (A, B)** Summary data quantifying ileal PCs **(A)** and IgA^+^, IgG^+^ and IgM^+^ PCs **(B)** for each of the three groups of study participants (PWVH, PWSH and NC). **(C, D)** Summary data quantifying colonic **(C)** and ileal **(D)** short-lived (SL), long-lived (LL) and very long-lived (VLL) PCs for each of the three groups. **(E, F)** Summary data quantifying colonic **(E)** and ileal **(F)** IgA^+^, IgG^+^ and IgM^+^ SL (left panels), LL (middle panels) and VLL (right panels) PCs for each of the three groups of study participants. Comparisons between groups were. performed using Kruskal-Wallis test with Dunn’s correction for multiple comparisons. Median and interquartile range was used to plot summary data, p values are shown as indicated.

**Figure S9**: **(A, B)** Manifold Approximation and Projection (UMAP) showing individual Ig gene distribution across PCs in the colon **(A)** and ileum **(B). (C)** Venn diagrams showing the numbers of shared and unique pathways found up-regulated (top panels) or down-regulated (bottom panels) pathways in the scRNA-seq and bulk RNA-seq datasets for each group’s comparison (PWVH vs NC; PWSH vs NC and PWVH vs PWSH). Comparisons were calculated using Fisher's Exact Test, p values as indicated.

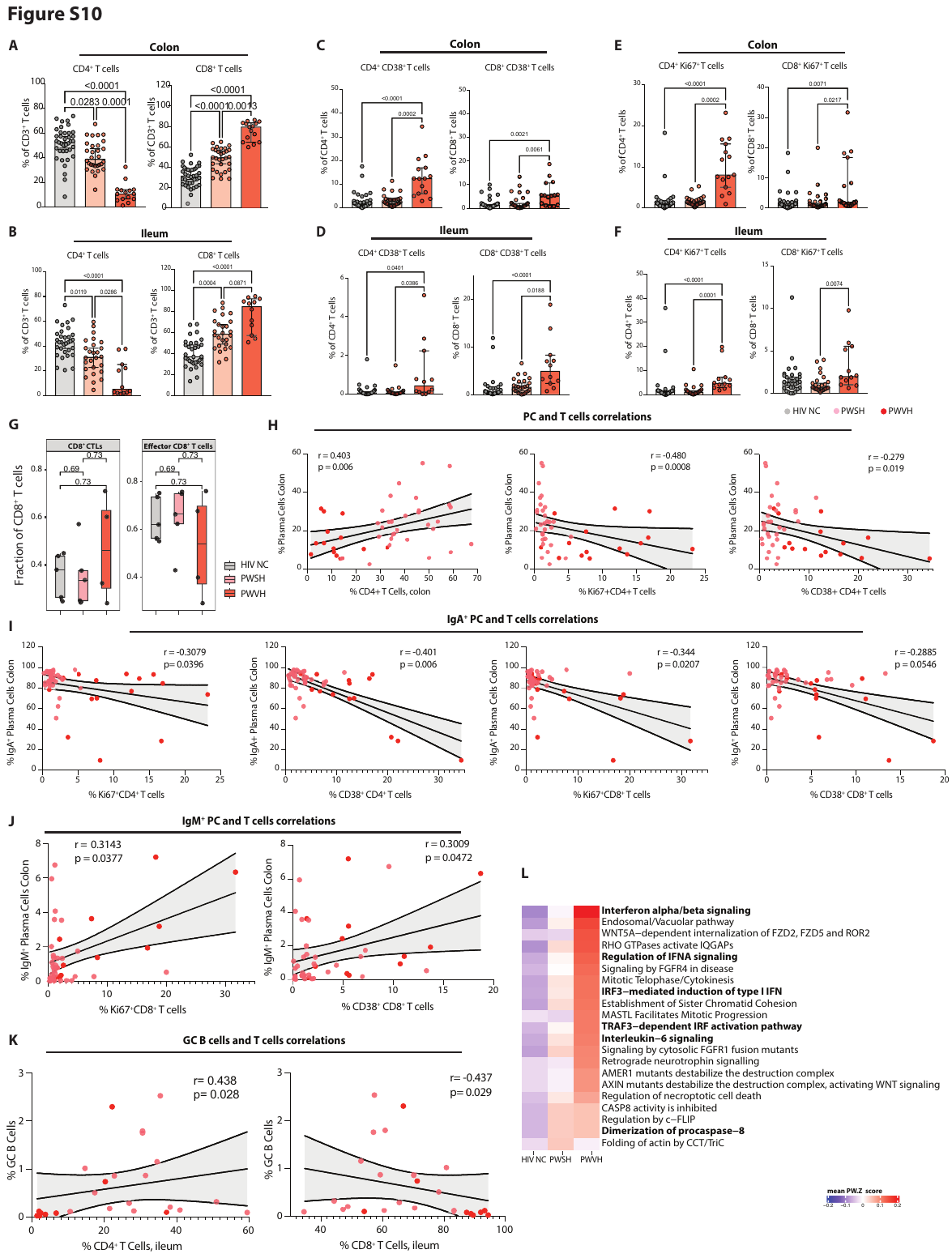

**Figure S10: Alterations in frequency and transcriptional profile of intestinal T cell subsets in PWH. (A, B)** Barplots comparing the frequency of colon- **(A)** and ileum-derived **(B)** CD4^+^ and CD8^+^ T cells across the three groups (PWVH, PWSH and NC) **(C, D)** Barplots comparing the frequency of colon- **(C)** and ileum-derived **(D)** activated (CD38^+^) CD4^+^ and CD8^+^ T cells across the three groups of study participants. **(E, F)** Barplots comparing the frequency of colon- **(E)** and ileum-derived **(F)** cycling (Ki67^+^) CD4^+^ and CD8^+^ T cells across the three groups. For panels A - F data is derived from the flow-cytometry dataset: (NC, n=51; PWSH, n=35; PWVH, n=16). Comparisons between groups performed using Kruskal-Wallis test with Dunn’s correction for multiple comparisons. Median and interquartile range was used to plot summary data, p-values are as indicated. **(G)** Boxplots comparing the frequency of ileum-associated CD8^+^ T cell subsets. Data is derived from scRNA-seq analyses. Comparisons performed with Wilcoxon test, p-values as indicated **(H-J)** Spearman’s correlation plot of colonic total CD4^+^ T cells, proliferating (Ki67^+^) and activated (CD38^+^) CD4^+^ T and CD8^+^ T cells with PCs (H), IgA^+^ PCs (I) and IgM^+^ Plasma Cells (J). (K) Spearman’s correlation plot of ileal T cells with Germinal Center (GC) B Cells. **(L)** Heatmap representing the mean z-score enrichment for selected pathways within the colonic TFH-like T cell subcluster.

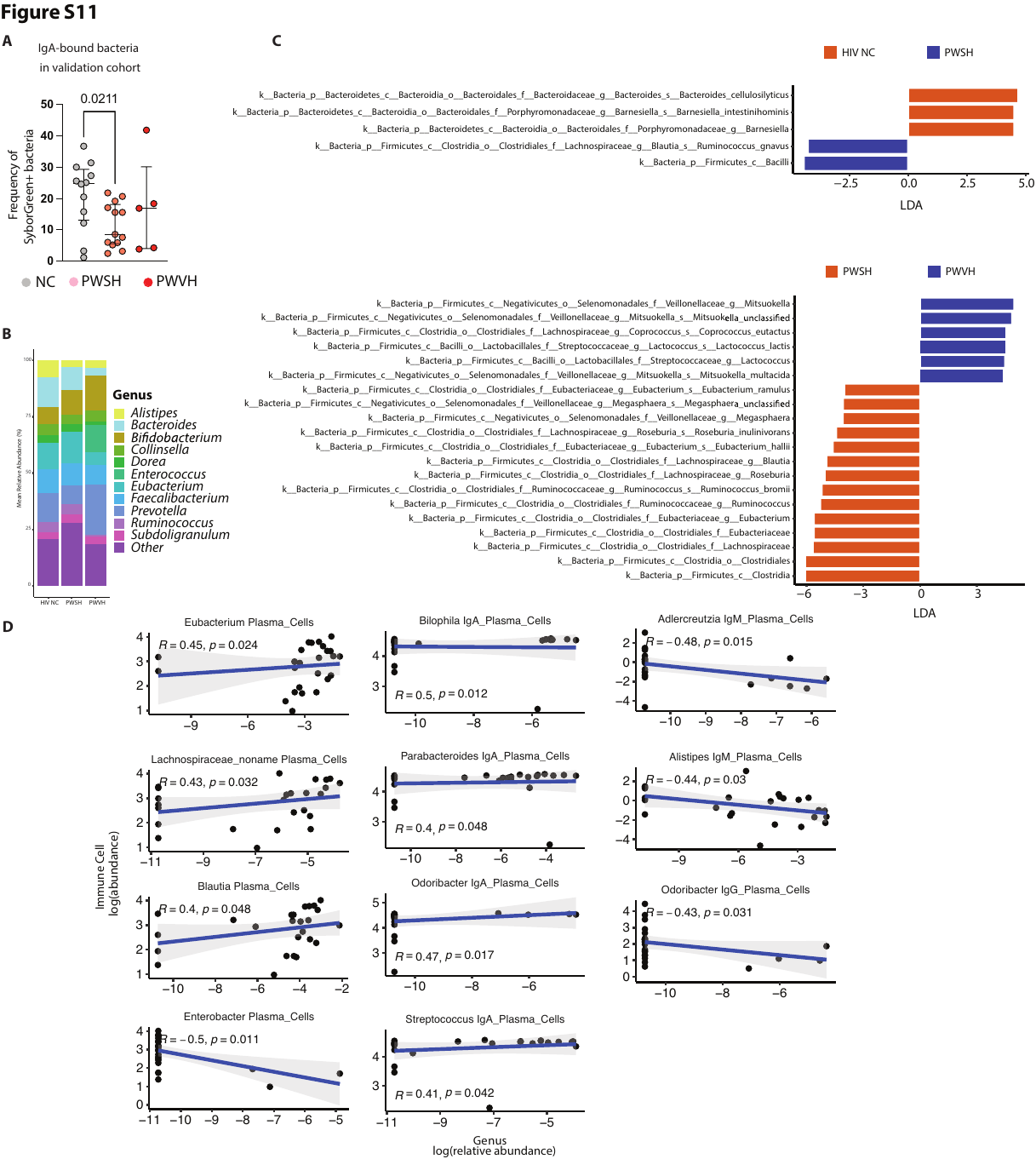

**Figure S11: Intestinal microbiota changes and targeting by secretory IgAs during HIV infection relate to altered colonic PCs Isotype expression. (A)** Summary data representing the frequency of IgA-bound bacteria across the three groups where stool samples were available for stool bacteria sorting in the validation cohort (Consortium for the Evaluation and Performance of HIV Incidence Assays (CEPHIA) of the University of California, San Francisco; 5 PWVH, 13 PWSH and 12 NC) **(B)** Barplot showing the distribution of bacteria genera across the three groups of study participants (NC, n=16; PWSH, n=18 and PWVH, n=7). **(C)** Linear discriminant analysis (LDA) effect size (LEfSe) performed on all Operational Taxonomic Units (OTUs) comparing PWSH and NC (top panel) and comparing PWSH and PWVH (bottom panel). **(D)** Correlation plots between the OTU-based relative abundance of bacterial genera and the frequency of colonic PCs and IgA^+^ PCs where a significant correlation was found using Spearman correlation. Unadjusted p-values as indicated.
